# Single-Cell Transcriptomic Attributes and Unbiased Computational Modeling for the Prediction of Immunomodulatory Potency of Mesenchymal Stromal Cells

**DOI:** 10.1101/2020.09.12.294850

**Authors:** Pallab Pradhan, Paramita Chatterjee, Hazel Y. Stevens, Chad Glen, Camila Medrano-Trochez, Angela Jimenez, Linda Kippner, Wen Jun Seeto, Ye Li, Greg Gibson, Joanne Kurtzberg, Theresa Kontanchek, Carolyn Yeago, Krishnendu Roy

## Abstract

Mesenchymal stromal cells (MSCs) are currently being tested in numerous clinical trials as potential cell therapies for the treatment of various diseases and due to their potential immunomodulatory, pro-angiogenic, and regenerative properties. However, variabilities in tissue sources, donors, and manufacturing processes and the lack of defined critical quality attributes (CQAs) and clinically relevant mechanism of action (MoA) pose significant challenges to identify MSC cell therapy products with a predictable therapeutic outcome. This also hinders regulatory considerations and broad clinical translation of MSCs. MSC products are often administered to the patient immediately after thawing from cryopreserved vials (out-of-thaw). However, the qualifying quality-control assays are either performed before cryopreservation, or after culturing the post-thaw cells for 24-48 hours (culture-rescued), none of which represent the out-of-thaw product administered to patients. In this study, we performed a broad functional characterization of out-of-thaw and culture-rescue MSCs from bone marrow (BM-MSCs) and cord tissue (CT-MSCs) using macrophage activation and T cell proliferation-based *in vitro* potency assays and deep phenotypic characterization using single-cell RNA-sequencing. Using this data, we developed unbiased computational models, specifically symbolic regression (SR) and canonical correlation analysis (CCA) models to predict the immunomodulatory potency of MSCs. Overall, our results suggest that manufacturing conditions (OOT vs. CR) have a strong effect on MSC-function on MSC interactions with macrophages and T cells. Furthermore, single-cell RNA-seq analyses of out-of-thaw BM and CT-MSCs indicate a tissue of origin-dependent variability and heterogeneity in the transcriptome profile. Using symbolic regression modeling we identified specific single-cell transcriptomic attributes of MSCs that predict their immunomodulatory potency. In addition, CCA modeling predicted MSC donors with high or low immunomodulatory potency from their transcriptome profiles. Taken together, our results provide a broad framework for identifying predictive CQAs of MSCs that could ultimately help in better understanding of their MOAs and improved reproducibility and manufacturing control of MSCs.

## INTRODUCTION

Mesenchymal stromal cells (MSCs) have been shown to be multipotent and immunomodulatory *in vitro*, and may have immunomodulatory, pro-angiogenic, and regenerative capabilities in vivo (1–3). Numerous clinical trials are currently underway where MSCs are being investigated as potential cell therapies to treat various diseases and immune disorders and mitigate acute or chronic inflammatory conditions. MSCs can be isolated from diverse tissue sources, e.g. bone marrow, cord tissue, and adipose tissue, and manipulated ex vivo for either autologous or allogeneic use (4–7). Allogeneic uses of MSCs are increasingly gaining popularity due to their off-the-shelf nature and ability to treat many patients from a single healthy donor. However, variabilities in MSC properties based on tissue-of-origin, donors, and manufacturing processes, and the lack of defined critical quality attributes (CQAs) to ensure functionally-predictive product comparability, make it difficult to select the best MSC product for clinical use (2, 5).

In a clinical setting, MSC-based cell products are commonly injected into the patients directly after thawing them from cryopreserved vials (3, 8) (described as out-of-thaw, OOT, hereafter). This is a more convenient approach than the logistically challenging options of using freshly harvested MSCs (before freezing) or culture-rescued MSCs where the cells are cultured *in vitro* for 24-48 hours after thawing from cryopreserved vials and before administration into patients (described as culture-rescue, CR, hereafter). CR cells require an on-site cell culture lab and technical personnel trained in aseptic technique. The cellular and molecular assays that are used for the assessment of MSC functions/potencies and often serve as release criteria, are typically performed on freshly harvested or culture rested/rescued MSCs rather than on straight out-of-thaw MSCs. CR cells, which are cultured for a couple of days in a cell culture plate or flask before running any functional/potency assays, are typically in a different metabolic state than the out-of-thaw MSCs and thus they may have different functions/potencies(8–10). Hence, it is imperative to understand the functional/potency differences between OOT and CR MSCs.

For the evaluation of immunomodulatory functions of MSCs *in vitro*, various *in vitro* assays using different types of immune cells and inflammatory conditions have been reported in the literature (2, 11, 12). Most of these *in vitro* functional assays are chosen based on hypothesized mechanisms of action (MoA) which are yet to be confirmed in relevant vivo models. T cell proliferation assessment is one of the widely reported assays which is often used for assessing the immunomodulatory functions of MSCs (11, 13) and as a potency assay to release MSC cell therapy products (12). However, recent mechanistic studies suggest that upon intravenous administration MSCs interact with phagocytic immune cells, such as macrophages/monocytes, which could then, through secretory factors or cell-to-cell interaction, influence other immune cells, including T cells, to mitigate inflammation or cause immunosuppression in vivo (14–16). In this context, macrophage/monocyte-based potency assays appear to be mechanistically more relevant for the evaluation immunomodulatory functions of MSCs.

Inherent variabilities in MSC cell therapy products due to differences in donors, sources and culture conditions make it difficult to identify their quality attributes that are predictive of their potency. There is an obvious need for deep characterization of MSCs from different donors, sources and culture conditions not only to understand the compositional differences between MSC products but also to find the signature attributes that correlate with their functions. Single-cell RNA sequencing of cells offer deep understanding of transcriptome profile of cells at single-cell level. Recently, we used single-cell RNA-sequencing (scRNAseq) to identify tissue of origin and cell cycle driven differential transcriptome profiles between bone marrow and cord tissue-derived MSCs (17). Here we report comparative functional characterizations of out-of-thaw and culture-rescue MSCs using macrophage and T cell-based immunomodulatory assays and use single-cell transcriptomic analysis coupled with unbiased computational modeling to identify transcriptomic attributes which are predictive of their immunomodulatory potency in macrophage and T cell assays.

## METHODS

### MSC expansion, freezing and thawing

Human bone marrow-derived mesenchymal stromal cells (BM-MSCs) were purchased from Roosterbio Inc. (Frederick, MD) at passage 2 (P2) and were expanded as per manufacturer’s instructions in T225 flasks for one passage to passage 3 (P3) using media supplemented with bovine serum (RoosterNourish-TM; described as regular media hereafter) or xeno-free additives (RoosterNourish-MSC-XF, described as xeno-free media hereafter). At >80% confluent, cells were fed fresh media, and harvested 24 hrs. later with TryPLE addition and subsequent washes with respective media (at harvest population doubling level, PDL 13-15). Umbilical cord tissue-derived mesenchymal stromal cells (CT-MSCs) were obtained from Duke University School of Medicine at passage 2 (P2) and expanded in Hyperflask culture (1720 cm^2^ total; 10 layers each with 175 cm^2^) for 5-7 days in Xeno Serum Free Media(XSFM) media (Prime-XV, Irvine Scientific, Santa Ana, CA) to passage 3 (P3). Cell pellets were resuspended in freezing medium (CryoStor5 for BM-MSC) or CryoStor10 for CT-MSC, Sigma-Aldrich, Saint Louis, MO) at a concentration of 5–10E6 cells/mL and aliquoted into vials. Cells were placed into a freezing container (Nalgene Mr. Frosty, Sigma, USA) in a −80°C freezer, to give a cooling rate of 1°C/min. After 24 hrs., frozen vials were transferred into storage in liquid nitrogen vapor phase. For thawing, vials were warmed in a 37°C water bath for 2 min and immediately transferred into diluting buffer (48 mL Plasmalyte A (Baxter, Deerfield, Illinois): 2mL human serum albumin (HSA 25%; Grifols albutein, Los Angeles, CA; final conc 1% HSA), the cell count and viability were determined on a NucleoCounter (Chemometec, Allerod, Denmark) before centrifuging at 500g for 10 min 4°C. Post centrifugation, cells were resuspended in diluting buffer to 2.5E6 cells/mL, recounted, and used in assays below

### THP-1 macrophage activation assay

THP-1 cells (acute monocytic leukemia TIB-202™; ATCC^®^, Manassas, VA) were thawed and seeded into 3 T75 suspension culture flasks at a density of 10E5/mL in RPMI-1640 (ATCC) media, supplemented with 20% fetal bovine serum (FBS, Hyclone heat inactivated (H.I.)) and 0.05 mM β-mercaptoethanol (BME; THP-1 expansion media), and expanded tenfold over 7 days. After splitting 1:5 (i.e. 15 flasks), by harvesting media from each flask and centrifuging at 300 g, 5 min, 4°C in a 50 ml conical tube, a second expansion was performed to ≤ 10E6/mL before harvesting and cryopreservation. Cell pellets were introduced gently into freezing media containing 10% DMSO: 90% FBS H.I. and after 24 hrs at −80°C in a freezing container, stored in vapor phase liquid nitrogen.

At point of assay, THP-1 cells were thawed in expansion media, counted, centrifuged at 300g, 5 min, 4°C and resuspended at 7E5/mL in expansion media for 3 days (6 mL in each of 2 T25 suspension flasks standing upright in incubator). After this time, the monocytes were harvested and the pellet resuspended in differentiation media (RPMI-1640 supplemented with 1% FBS H.I., Penicillin/Streptomycin (P/S, VWR, Radnor, PA) and 100 ng/mL phorbol 12-myristate 13-acetate (PMA; Sigma-Aldrich). No BME was added to the media. The cells were plated at 100,000/well in a CELLBIND 96 well plate and left to adhere 48 hrs. PMA treatment promoted the differentiation of monocytes to naïve macrophages. On the same day, culture rescued MSCs were set up, using the thawing technique of diluting buffer for MSCs, and then plating at 3700 cells/cm^2^ in a T225 flask in the appropriate growth media for BMMSCs or CTMSCs. Culture rescue was for 48 hrs at 5% CO_2_, 37°C.

Culture rescue MSC were harvested by TryPLE addition and washed with growth media. Cells were counted pre-spin, centrifuged at 500g/10 min 4°C before resuspending in diluting buffer at 2.5E6 cells/mL and recounted. Out of thaw MSC (OOT) were prepared by diluting freshly thawed cells in diluting buffer, centrifuging 500g/10 min 4°C and then concentrating to 2.5E6/mL and recounting. Both pre-spin and post-spin cell counts, and viability were determined. The naïve macrophage plate was spun at 300g/5min at 4°C to settle any loosely adherent cells before changing media. Media was aspirated, taking care to avoid removing cells. 180 uL of co-culture media (RPMI-1640 media containing 1% FBS, LPS (100 ng/mL; from E coli, Sigma-Aldrich) and IFNɣ (100 ng/mL; PeproTech, Rocky Hill, NJ) (Pietila, M) was added to 20 uL cell suspension of CR or OOT MSCs (100,000 cells to give THP-1M:MSC, 1:1) and immediately added to each well (n=6). Co-culture media was also added to separate wells containing THP-1M or MSC only. The plate was incubated for 24 hrs at 5% CO_2_, 37°C. After this time the plate was centrifuged at 500g/5 min 4°C before removal of conditioned media to a fresh 96 well plate for storage at −20°C until time of assay.

### ELISA and Luminex assay for growth factors/cytokines

Cell culture supernatant (diluted ¼) was subjected to ELISA (TNFα, Thermo Scientific, Waltham, MA, n=6) and Luminex (Cytokine Human Magnetic 30 Plex Panel, Thermo Scientific, n=4) per manufacturer’s instructions. Results from ELISA are indicated as TNF-α concentration (pg/mL) and % inhibition of TNF-α released by THP-1M alone. Results from Luminex are shown in a Z score heatmap.

### T cell proliferation assay

The immunomodulatory effect of MSC on T cell proliferation was evaluated using *in vitro* cocultures between MSCs and PBMCs. In brief, MSCs were co-cultured with CFSE-labeled, Anti-CD3/CD28 bead (Dynabeads)-activated PBMCs (MSC/PBMC at 1:2) from healthy volunteers (purchased from Zen-Bio) plated in 200 ul culture media (RPMI 1640 + 10% FBS + 1% PS+ 30 IU/mL IL2) in a 96-well plate for 72 hrs. Thereafter, the cells were harvested and stained with CD4 and CD8 T cells using anti human CD3, CD4, CD8 and CD25 antibodies for flow cytometry and analyzed using a BD-Fortessa flow cytometer. T cell proliferation level was assessed by quantifying CFSE dilution using FlowJo software. Secreted factors were measured in the T cell assay culture media using a Luminex kit (Cytokine Human Magnetic 30 Plex Panel, Thermo Scientific) per manufacturer’s recommendations with 4 replicates per sample. Observed concentration values of samples were obtained in reference to standard curves run on each plate. Concentration values were analyzed for outliers using Grubbs’ statistical analysis with alpha of 0.05 for each donor in respective group (i.e., OOT, CR). Mean values were calculated for each group and normalized with Z score using JMP statistical software. Z scores were used for creation of heatmap.

### Single-cell RNA sequencing sample preparation and data analysis

We used OOT MSCs from 7 BM-MSC (P3) and 3 CT-MSC samples (P3) for single-cell RNA sequencing. Briefly, cryopreserved MSC vials were thawed for 2 minutes at 37° C in a water bath and mixed cold Plasmalyte + 1% HSA and centrifuged at 200g for 5 minutes. Next, the supernatant was removed, and the cell pellets were resuspended, and cell viability and counts were performed.

The cells were then filtered and counted to adjust the volume and concentration. The filtered cells were maintained on ice until next step. The cells were counted using a TC20 system and centrifuged briefly and re-suspended in an appropriate volume of cold PBS containing 0.1% BSA to achieve a final concentration of at least 2500-2700 cells/μl.

Next, the single cell suspension was loaded onto a ddSEQ cartridge, and the cells were encapsulated and barcoded using a single-cell Isolator. Lysis and barcoding took place in each droplet. Droplets were disrupted and cDNA was pooled for second strand synthesis. Libraries were generated with direct cDNA tagmentation using Nextera technology. Tagmentation was followed by 3ʹenrichment and sample indexing to prepare indexed, sequencing-ready libraries. The libraries were sequenced using Nextseq500 sequencing Platform (PE75, mid-output V2.5 kit). Upon collecting data, Basespace Surecell RNA single cell software was used to demultiplex and align the samples and generate expression matrix. The downstream analysis was done using SC3 and SC-pool pipelines(17). EdgeR was used to extract differentially expressed genes from the single-cell RNA-seq data. The downstream analysis was done using SC3 (18) for the clustering analysis and Seurat (19) for normalization, scaling, PCA and differential expression analysis, and Wilcoxon Rank Sum test was implemented in Seurat (20).

### Statistical analysis

Statistical analysis and all graphs were performed in Prism 8.4 except the principal component analysis (PCA) which was done in Python. All data are presented as mean ± standard deviation (SD) unless otherwise indicated. The significance level was determined by a one-way analysis of variance (ANOVA) with post-hoc Tukey’s comparisons for the data with normal distribution and by nonparametric Kruskal-Wallis with post-hoc Dunn’s comparison for the data with non-normal distribution. Shapiro-Wilk rest was performed to test normality. A p value <0.05 was considered significant.

### Predictive computational modeling

We used symbolic regression (SR) to identify the single-cell transcriptomic attributes of BM and CT-MSCs that predict the immunomodulatory potency in THP1-macrophage activation and T cell proliferation assays. For SR, the top 1000 differentially expressed genes between BM and CT-MSCs were allowed as potential inputs for predicting the targeted responses, i.e., percentage of TNF alpha suppression (normalized with THP-1 macrophage only group) or percentage of CD4 T cell proliferation (normalized with activated PBMC only group) for OOT and CR conditions. We used DataModeler software (Evolved Analytics) for symbolic regression modeling.

Canonical correlation analysis (CCA) was applied to the same set of genes (inputs) and potency assays (outputs) to create four separate models. In each model, a subset of donors was used to train and test the model (50/50 train-test split) with the remaining donors used as validation of predictive capacity. The input data was scaled using a quantile transformation to minimize the influence of outliers and the output data was scaled to a range of (0-1). The scaling and model transforms were fit using the training set and equivalently applied to the remaining test and validation data. CCA analysis was performed using Python2.7.

## RESULTS

We used 7 bone marrow and 3 cord tissue derived MSC samples (**Table S1**) for functional and omics characterization (see workflow in **Figure 1**). For bone marrow-derived MSCs (BM-MSCs), we used 5 donors expanded with serum supplemented regular media and 2 matching donors expanded with xeno-free media. All the BM-MSCs cells were purchased from RoosterBio at passage 2 (P2) and expanded one passage further (P3) using RoosterBio expansion protocol, and all 3 cord-tissue derived MSCs (CT-MSCs) at passage 2 (P2) were obtained from Duke School of Medicine and expanded up to passage 3 (P3) using Duke’s expansion protocol and cryopreserved before using them for any functional and omics characterization. For functional characterization, we used macrophage activation and T cell proliferation assays to comparatively evaluate the immunomodulatory effect of out-of-thaw (OOT) and culture-rescue (CR) BM- and CT-MSCs. For OOT condition, cryopreserved vials are thawed and directly used for the characterization. For CR condition, cryopreserved vials are thawed, seeded in the tissue culture plates or flasks and cultured for 48 hours before any functional characterization. For omics characterization, we employed single-cell RNA-sequencing for BM and CT-MSCs for OOT samples only and considered the OOT MSCs as cell therapy product. Next, we developed predictive modeling based on symbolic regression (SR) and canonical correlation analysis (CCA) using single-cell transcriptome as “input/predictor” and Macrophage and T cell potency assay results as “outputs/responses” to identify attributes and donors that are predictive of the immunomodulatory potency of MSCs.

**Figure 1.**
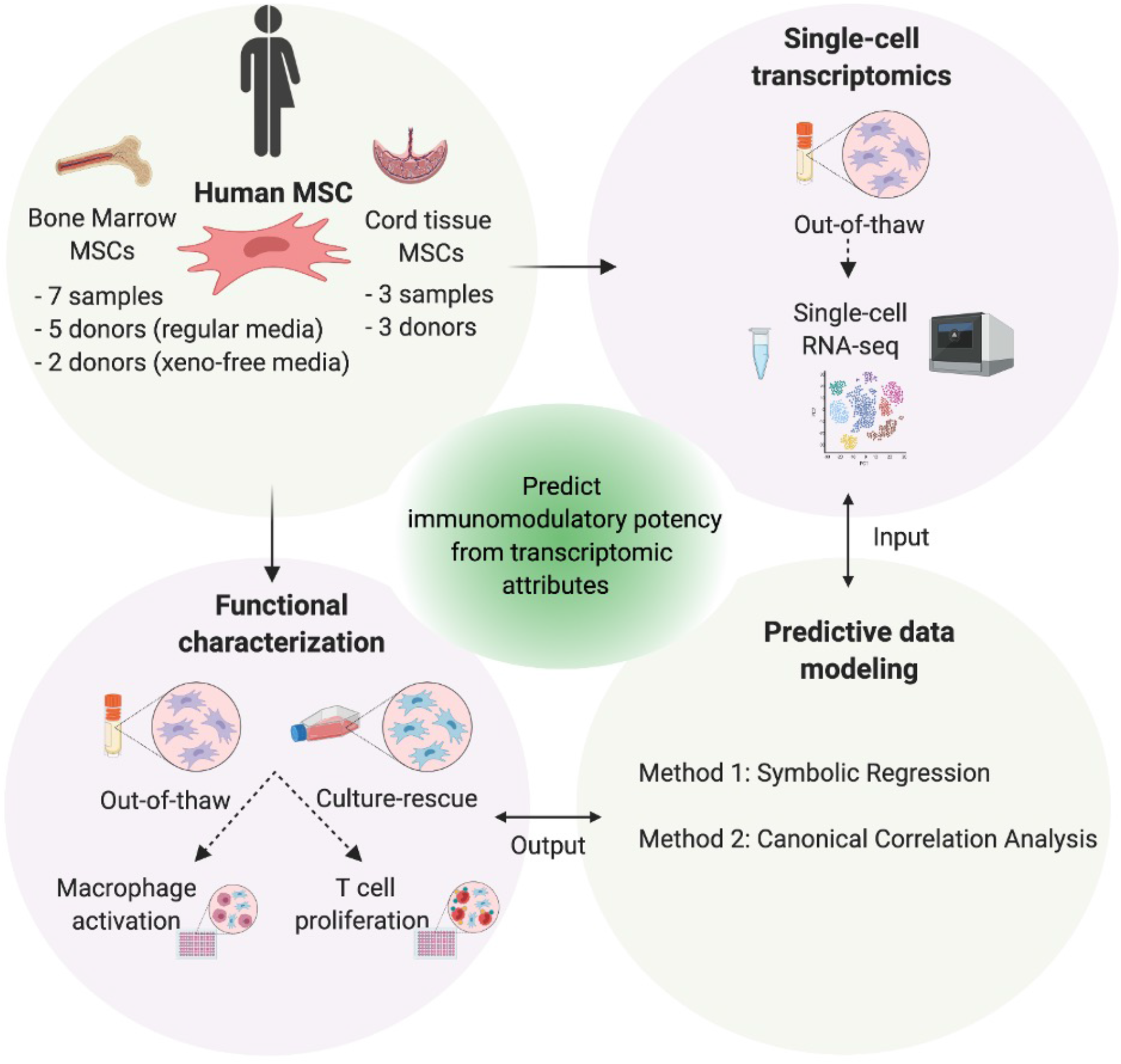
Schematic illustration showing workflow for functional and single-cell transcriptomic characterization of MSC and identification of predictive attributes through predictive data modeling.

### Both out-of-thaw and culture-rescue MSCs suppress proinflammatory cytokines from activated THP-1 monocyte-derived macrophages

We used the THP-1 macrophage activation assay to evaluate the immunomodulatory effect of BM and CT-MSCs for OOT and CR conditions. In this assay, we stimulated THP-1 monocyte-derived macrophage with LPS and IFN-ɣ for 24 hrs and quantified secreted factors (including cytokines, chemokine and growth factors) using a 30-plex Luminex kit and ELISA (for TNF-α only) (**Figure 2A**). Both BM and CT MSCs suppressed multiple cytokines and chemokines as compared to the THP-1 macrophage group which showed high levels of a number of pro-inflammatory cytokines and chemokines in this assay (**Figure 2B**). Moreover, principal component analysis (PCA) of the secretome shows clear separation between the THP-1 macrophage only group and THP-1 macrophage-MSC coculture group in the PC1 (32.8%) axis (**Figure 2C**), suggesting differential expression of secretome between these groups. Interestingly, OOT and CR conditions show some separation in the PC2 (16.6%) axis, indicating some differences in the secretome levels between these two conditions (**Figure 2C**). The separation between OOT and CR conditions was more apparent in the PCA plot when we excluded the THP1 macrophage group from the analysis (**Figure S1A**). Interestingly, multiple pro-inflammatory cytokines, such as TNF alpha, IL12, MIP-1α, MIP-1ß show high PC1 scores (**Figure S1B**). In addition, IL1-RA, an anti-inflammatory cytokine, also showed up as one of the top variables on PC1, and MIG (CXCL9) chemokine was one of the top contributors in PC2 variability (**Figure S1B**). When we individually inspected individually at these cytokines comparatively across different groups, we found that both OOT and CR conditions for BM and CT-MSC donors were able to suppress TNF alpha, IL12, MIP-1α, MIP-1ß, IL1-RA cytokines (**Figure 2D-H**). Interestingly, as indicated by the PC2 scores (**Figure 2C and S1A**), OOT and CR conditions show a differential response for MIG chemokine with only CR condition being suppressive (**Figure 2I**). Furthermore, we found some variability across donors and differences between OOT and CR conditions in the TNF alpha response for BM and CT-MSCs (**Figure S1C**).

**Figure 2.**
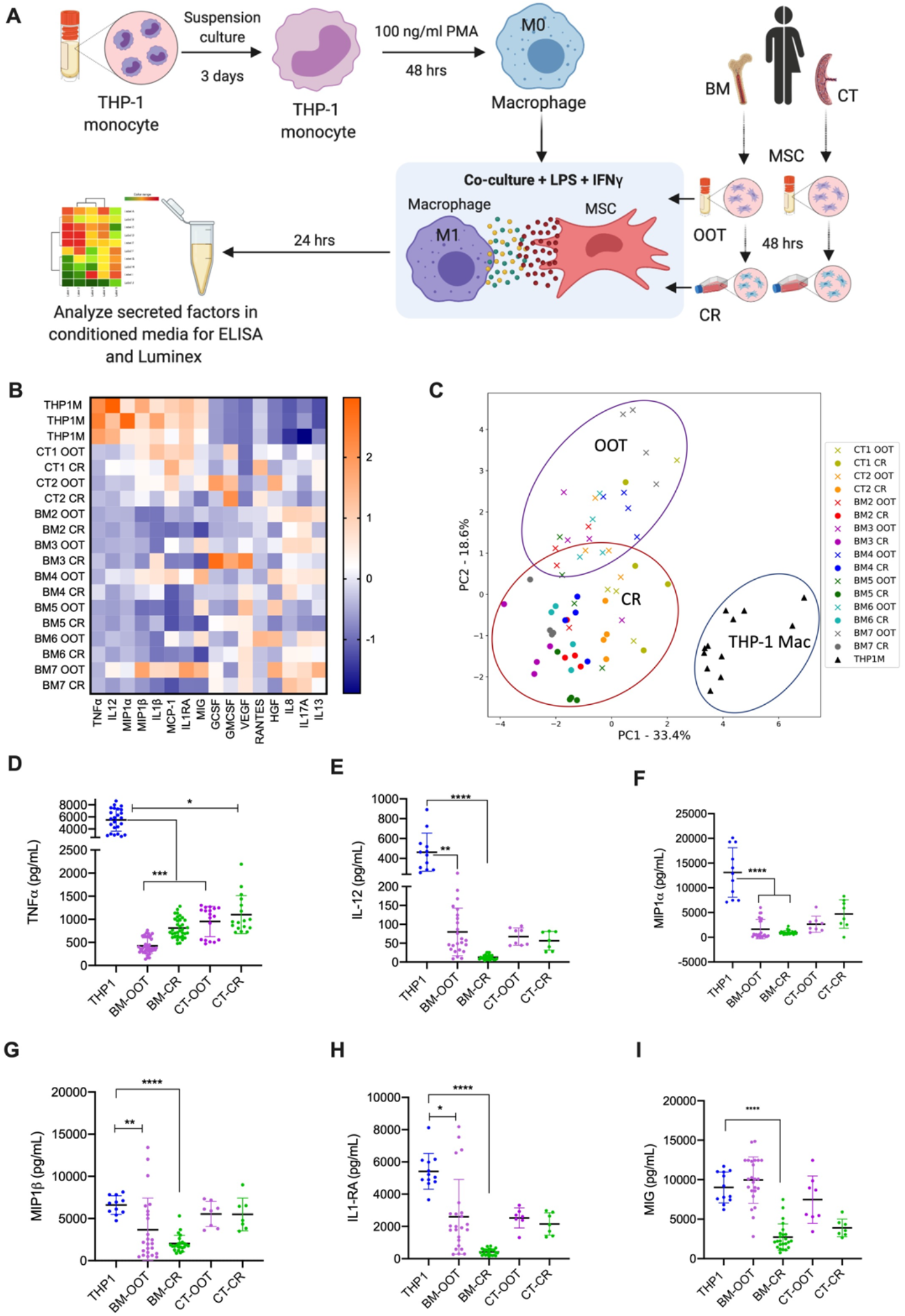
Immunomodulatory effect of BM- and CT-MSCs on cytokines, chemokines, and growth factors in THP-1macrophage activation assay. A) Macrophage activation assay workflow; B) Heatmap (average Z score) showing cytokine, chemokine and growth factor levels for BM and CT-MSCs for OOT and CR conditions; C) Principal component analysis (PCA) plot (PC1 vs PC2) of cytokine and chemokines levels for BM and CT-MSCs; Levels of cytokines and chemokines with high PC1 and PC2 loading scores in the PCA - D) TNF-α, E) IL-12, F) MIP-1α, G) MIP-1ß, H) IL1-RA and MIG (CXCL9). Data represent mean ±SD. Kruskal-Wallis test with Dunn’s post-hoc test was performed. **P<0.01. ***P < 0.001, ****P<0.0001.

### Out-of-thaw MSCs exhibit very low or no T cell suppression whereas culture-rescue MSCs show moderate to high T cell suppression across BM and CT-MSC donors

Next, we evaluated the immunomodulatory effect of BM and CT-MSCs on T cell proliferation for OOT and CR conditions using a T cell proliferation assay where MSCs were co-cultured with CFSE-labeled, anti-CD3/CD28 coated bead-activated PBMCs for 3 days before flow cytometry analysis in the media from T cell proliferation assay using a multiplex Luminex kit (**Figure 3A**). In this assay, OOT MSCs showed no or very low T cell proliferation suppression for most of the donors across BM and CT-MSCs (**Figure 3B and C: flow histogram in Figure S2A**). However, CR MSCs showed high T cell proliferation suppression with some degree of donor variability across BM-MSC and CT-MSC donors (**Figure 3B and C; Figure S2A**). Furthermore, we observed comparable T cell proliferation suppression between CR BM-MSCs from the same donor that were either cultured with regular or xeno-free media (BM2-xeno-free vs BM6-regular, and BM3-xeno-free vs BM7-regular, **Figure 3B**).

**Figure 3.**
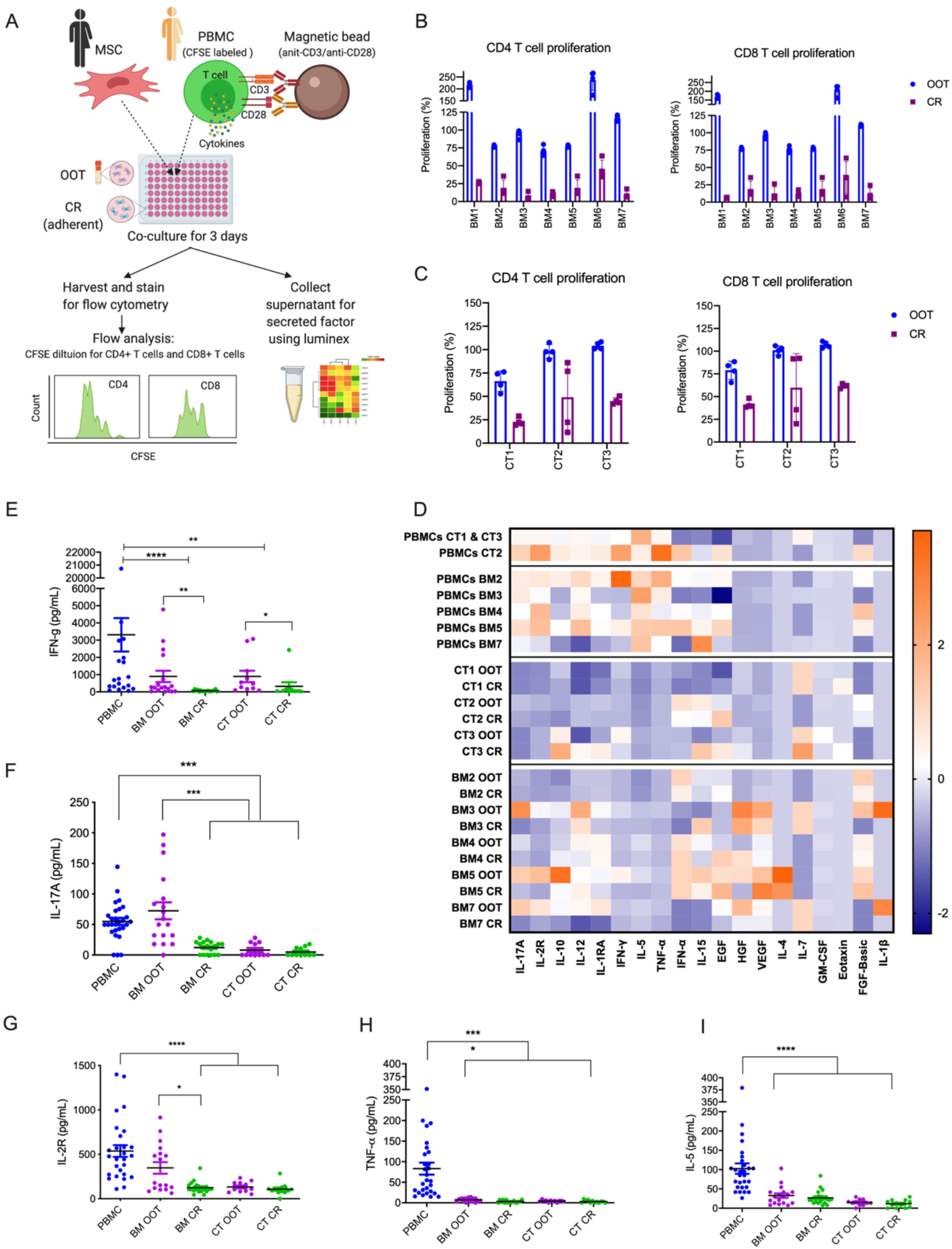
Immunomodulatory effect of BM and CT-MSCs on T cell proliferation for OOT and CR conditions. A) T cell proliferation assay workflow; Percentage of CD4 and CD8 T cell proliferation normalized to activated PBMC group (with 100% proliferation) for B) BM-MSC and C) CT-MSC donors; D) Z-score heatmap showing levels of secreted factors in T cell assay media; Comparative levels of individual cytokines - E) IFN-gamma, F) IL17-A, G) IL-2R, H) TNF-α, I) IL-5 cytokines in T cell assay media. Data represent mean ±SD. Kruskal-Wallis test with Dunn’s post-hoc test was performed. **P<0.01. ***P < 0.001, ****P<0.0001.

For CR condition in our T cell proliferation assay, we first seeded MSCs in a 96-well plate and cultured for 48 hours and then added PBMCs and T cell activation beads (Dynabeads) to the plate without harvesting the MSCs. However, for potential use of CR MSCs in clinics, MSCs need to be harvested following culture rescuing. Therefore, we compared the effect between adherent and harvested culture rescued MSCs on T cell proliferation. For harvested culture-rescue conditions, we seeded MSCs in both low density and high density (3700 or 50,000 cells/cm^2^) to evaluate the effect of seeding densities on T cell proliferation. Overall, adherent CR MSCs showed relatively higher T cell suppression as compared to the high- or low-density harvested CR and OOT conditions, and both high- and low-density CR conditions showed high variability in T cell proliferation suppression across MSC donors (**Figure S2B**). In addition, we used two PBMC donors to compare T cell proliferation suppression capacities of MSCs and observed similar results for both PBMC donors (**Figure S3**). Also, when we compared 3-day proliferation assay (our standard duration for T cell proliferation assay) with 5-day proliferation assay, we observed similar results for both durations for T cell proliferation by CSFE dilution **(Figure S4**).

Next, we measured secreted factors in the T cell proliferation assay media and found that both BM and CT MSCs had lower levels of multiple cytokines as compared to activated PBMC groups with some degree of variability between donors and culture conditions (OOT vs CR) **(Figure 3D).** Then, we compared some key T cell related cytokines across BM and CT donors for OOT and CR conditions. For IFN-gamma, CR conditions showed significantly lower levels than activated PBMCs and OOT conditions for both BM and CT-MSCs **(Figure 3E**). For IL-17 and IL-2R, CR condition for BM-MSCs and both OOT and CR conditions for CT-MSCs had significantly lower levels than activated PBMCs (**Figure 3F-G**). Furthermore, both OOT and CR conditions for BM and CT-MSCs showed significantly lower levels of TNF-α and IL-5 compared to activated PBMCs (**Figure 3H-I**).

### Single-cell RNA-seq analysis of BM and CT-MSCs indicate tissue source and culture condition-dependent differential gene expression

We performed single-cell RNA-sequencing using 10 samples - 7 BM-MSC and 3 CT-MSC samples. We included two BM-MSC donors that were cultured and expanded using both regular and xeno-free media. Details on the single-cell sequencing parameters and sample and donor information are shown in Table S2.

Even though they are all categorized as “MSCs” by the ISCT consensus surface markers, BM and CT-MSCs show different gene expression profiles. In **Figure 4A**, PC1 (52%) separates cells by their UMI counts, while PC2 (5%) separates cells by tissue sources for MSCs. MSC cultured in xeno-free media are located between bone marrow and cord tissue-derived MSCs. Cells with low UMI counts were not included in the rest of the analysis.

**Figure 4.**
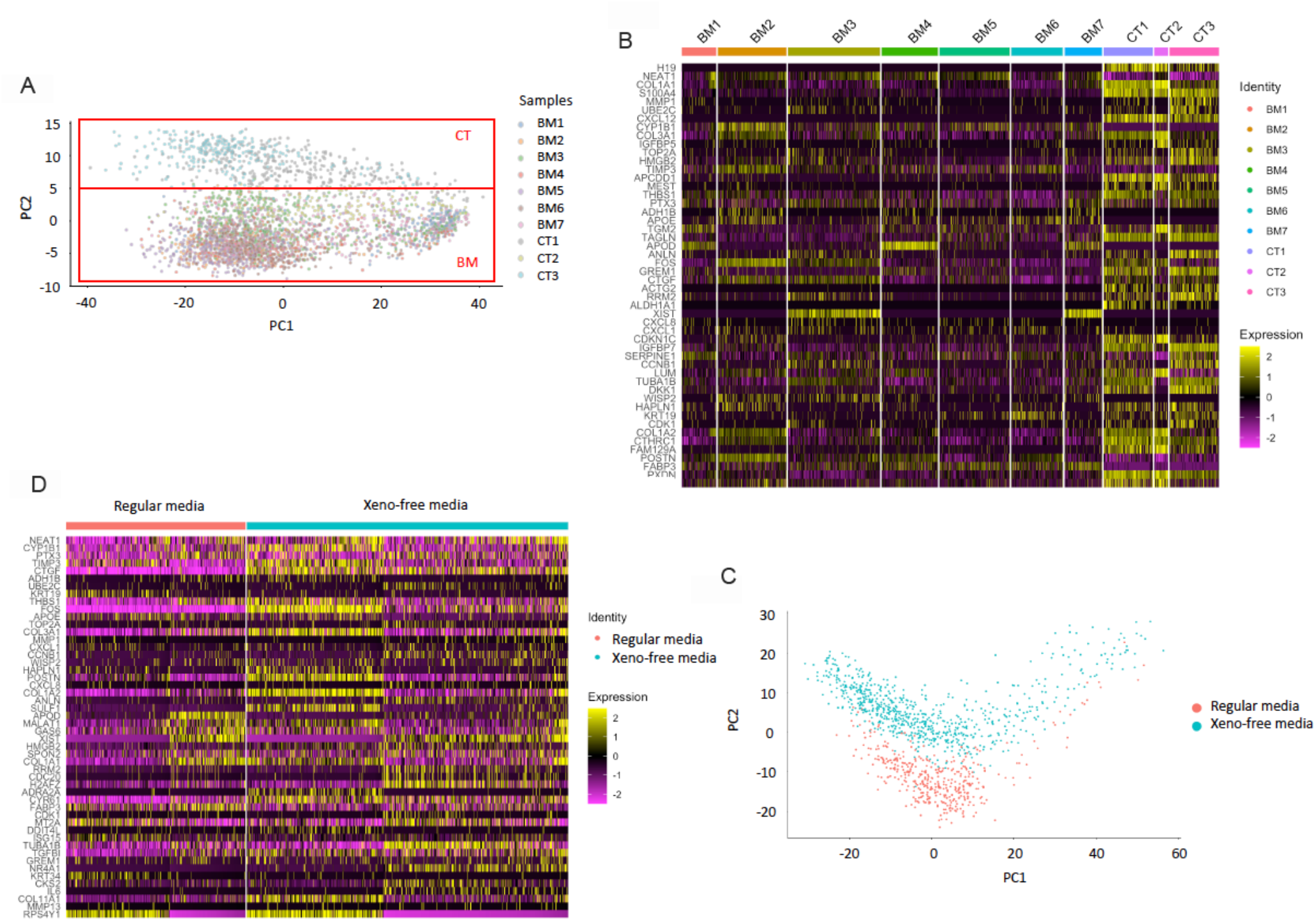
Single-cell RNA-seq analysis of BM and CT-MSCs. A) Gene expression PCA plot showing clustering of BM and CT-MSC samples; B) Heat map shows topmost differentially expressed genes between BM and CT-MSC samples; C) Gene expression PCA plots showing clusters for BM-MSCs expanded with regular and xeno-free media; D) Heat map Topmost differentially expressed genes between samples expanded in regular media and samples expanded in xeno-free media.

A differential expression analysis was performed between BM and CT-MSCs. More than 5000 genes were found to be differentially expressed between BM and CT-MSCs, at an FDR <5%. **Figure 4B** highlights the topmost variable genes between BM and CT-MSCs. Similar to our previous findings (17), both BM and CT-MSCs were divided in two major sub-clusters – BMa and BMb or CTa and CTb. For BM-MSCs, a differential expression analysis was done between these two clusters, followed by a gene ontology analysis showing cluster BMa overexpressed genes are involved in immune signaling and other pathways related to MSC functions, while cluster BMb are involved in mitotic cell cycle. We have reported a detailed analysis of BM and CT-MSC subcluster-dependent differential expression and gene ontology recently (17).

After determining the differences between BM and CT-MSC samples and the heterogeneity of gene expression within each tissue type, we looked at the differences in gene expression for BM-MSC samples based on the types of culture media (regular vs xeno-free media) used for expanding the MSCs from same donors. As shown in **Figure 4C**, regular and xeno-free media expanded BM-MSCs cluster separately in the PCA plots, suggesting culture media-dependent gene expression for BM-MSCs. **Figure 4D** shows top differentially expressed genes between regular and xeno-free media for BM-MSCs.

### Symbolic regression models identify single-cell transcriptomic attributes for immunomodulatory potency of MSCs

We used symbolic regression (SR) to identify single-cell transcriptomic attributes of BM and CT-MSCs that predict the immunomodulatory potencies of MSCs in the macrophage activation and T cell proliferation assays. For SR, we used the top 1000 differentially expressed genes (FDR < 5%) between BM and CT-MSCs (all OOT samples) were allowed as potential inputs for predicting the targeted macrophage activation or T cell proliferation results for OOT and CR conditions. We used percentage of TNF-α suppression normalized to THP1-macrophage only group and percentage of CD4 T cell proliferation normalized to activated PBMC only group as MSC immunomodulatory potency output parameters. We assigned the same potency “output” to the “input” transcriptome of every cell within each donor sample. The models were run under two conditions – a) allowing tissue types and all cells as potential variables and b) allowing all cells, but not allowing tissue type as a potential variable. The top performing models yielding the highest accuracy and lowest complexity were used to identify potential driving variables (**Table S3**) and variable combinations for each response (**Table S4-S7**). As shown in **Figure 5 and Table S3**, when tissue type is allowed as a potential variable, it emerges as a top driving variable for MacTNFa-OOT, MacTNFa-CR, and CD4Poliferation-CR, but not for CD4Poliferation-OOT responses. MacTNFa-OOT yields the highest R2 values, followed by MacTNFa-CR, CD4Poliferation-OOT and CD4Poliferation-CR (**Table S3**). Additionally, SR models identify a number of transcriptomic features as major driving variables even when the tissue types were allowed as a potential variable (**Figure 5** and **TableS3**).

**Figure 5.**
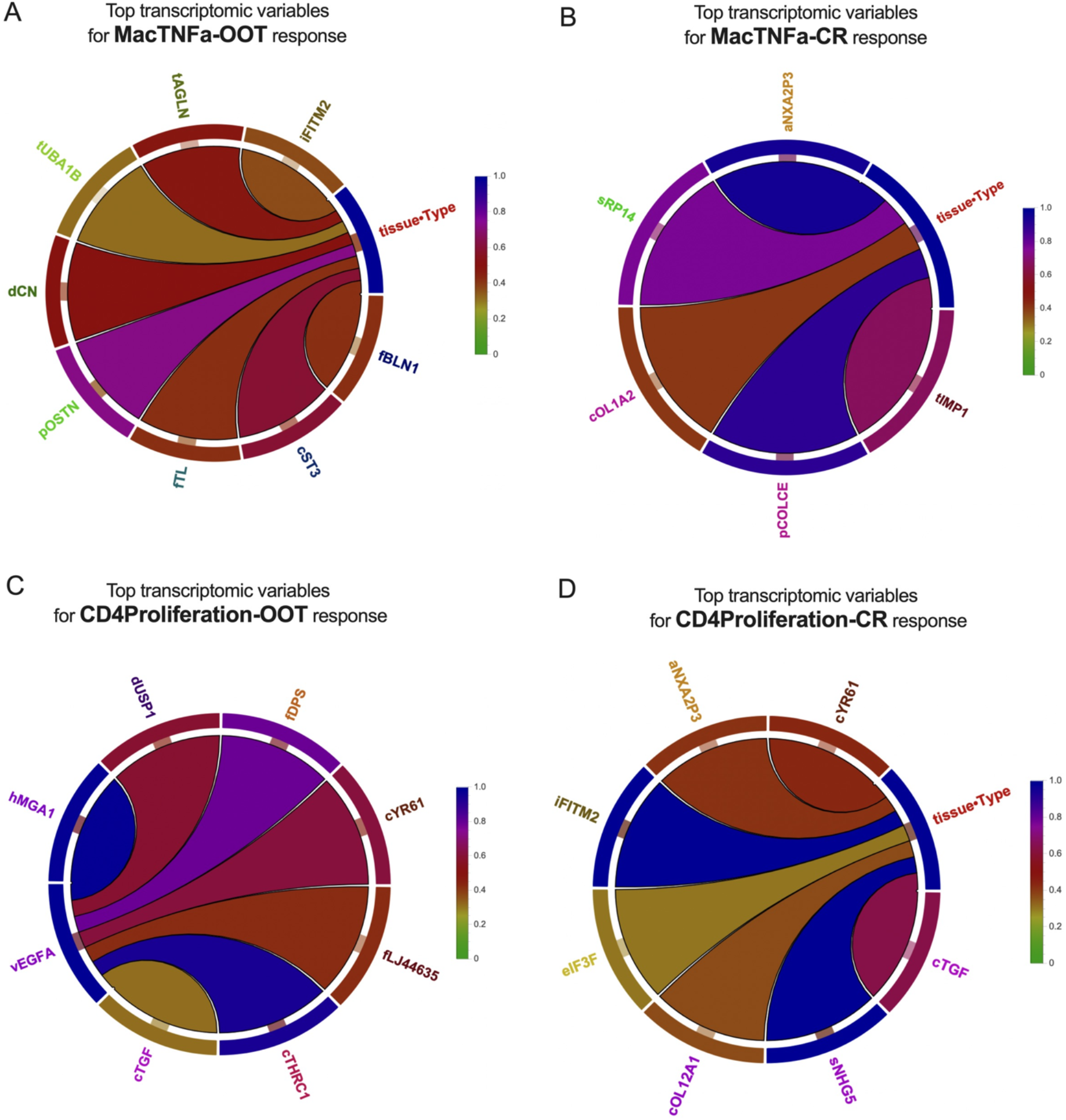
Symbolic regression (SR) models identify transcriptomic attributes that predict the immunomodulatory potency of MSCs. Variable association core diagrams show the top transcriptomic variables which are present in > 30% of the top performing models and the pairwise frequency of the most dominant variable for A) macrophage TNF-alpha response for OOT, B) macrophage TNF-alpha response for CR, C) CD4 T cell proliferation response for OOT, and D) CD4 T cell proliferation response for CR conditions for BM and CT-MSC samples. In these models, tissue source of MSCs in addition to all cells from each donor were allowed as potential variables. The colors of the connections indicate the frequency and strength of the variable association.

Importantly, the transcriptomic features that showed up as top variables mostly differ across potency responses/outputs. Only a few features, such as IFITM2, CTHRC1, PXDN, LTBP1, SERINC2 and DUSP1 were identified as common variables for more than one potency outputs; however, there was no common variable present across all four potency outputs (Table S3).

### Canonical Correlation Analysis models predict MSC donors with high or low immunomodulatory potency from their single-cell transcriptome profile

Next, we ran Canonical correlation analysis (CCA) to predict the potency of MSC donors (high vs low potency donor) from their single-cell transcriptome profiles. We developed CCA models using the top 1000 differentially expressed genes as “inputs” and applied to each output metric for macrophage TNF alpha suppression and CD4 T cell proliferation levels for OOT and CR conditions. For each model, a subset of donors was selected to train and test the model (shaded gray in the table in **Figure 6**). The remaining donors that were not used in the training of the model were used to validate the predictive capacity (shaded orange in the table in **Figure 6**). The first canonical correlation component (CC1) for each test and validation dataset was calculated and plotted as a density distribution (**Figure 6**). The CC1 represents a relative measure of potency by maximizing the correlation between the input scRNAseq data and the output metric (MacTNFa-OOT, MacTNFa-CR, CD4Proliferation-OOT, CD4Proliferation-CR). The relative potency of each cell sample (i.e. the target output metric) is depicted by the location of the distribution along the x-axis (arrows high vs low potency in **Figure 6**).

**Figure 6.**
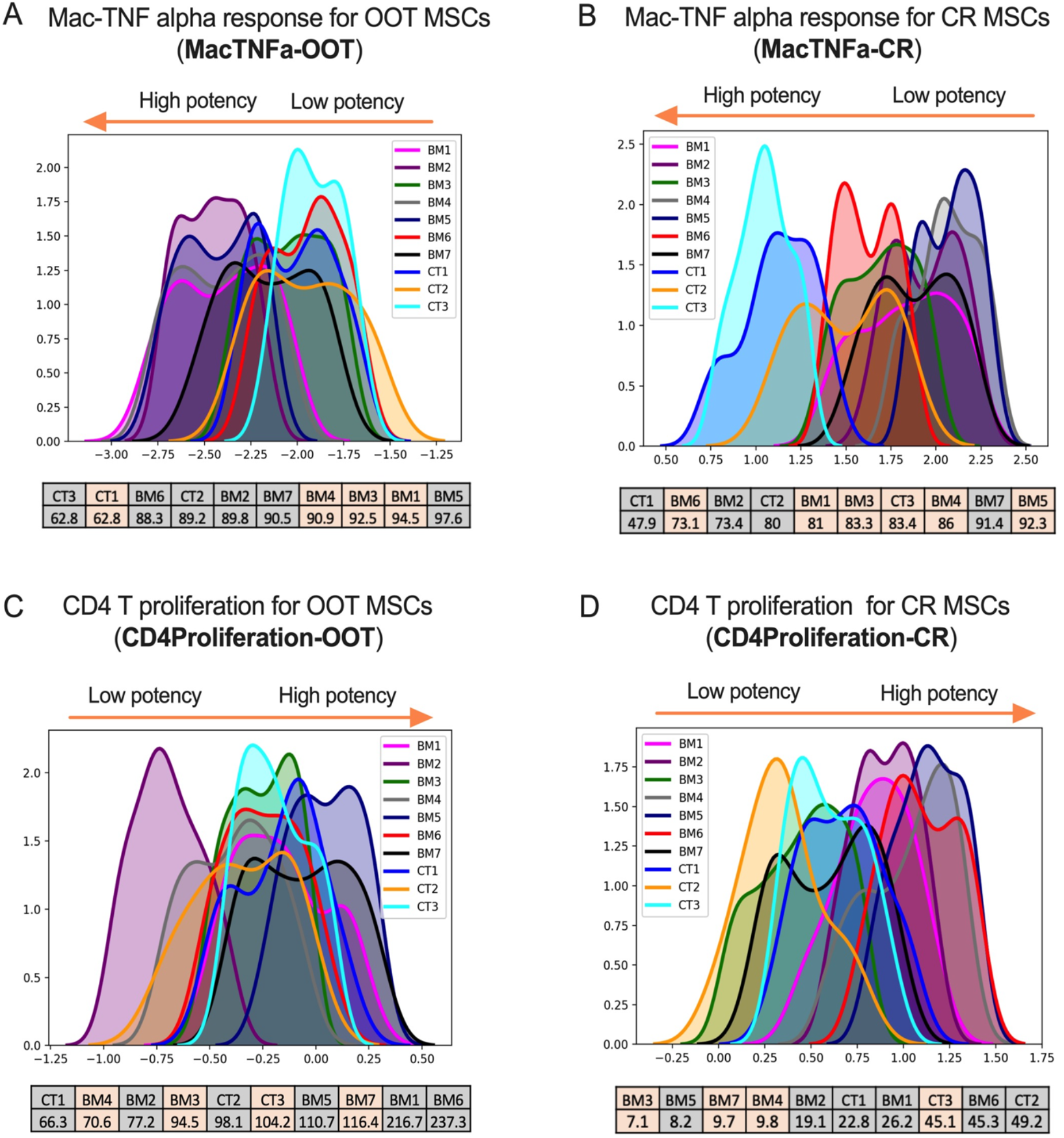
CCA models predict MSC donors for immunomodulatory potency from their single-cell transcriptomic features. Density distribution of first canonical correlation components (CC1) for macrophage TNF alpha responses for A) OOT and B) CR MSCs and CD4 T cell proliferation responses C) OOT and D) CR MSCs. The location of the distribution represents the maximum correlation between the normalized RNA-seq data and the respective output metric. Positive and negative values do not translate to positive or negative potency, but the relative order of the donors represents the rank of potency. Orange highlights in the table associated with each figure represent 100% predicted samples.

For the MacTNFa-OOT output (**Figure 6A),** the model identified three main transition points of low, medium, and high potency by the gene expression in donors CT3, BM7, and BM5, respectively. BM1 and BM4 donors were both correctly identified to have high potency and cluster with the high potency sample, BM5. CT1 and CT2 were also correctly predicted to cluster with the lower potency donors CT3 and BM6. The donor BM3 is the only sample that did not align with the true order of potency. However, a meta-analysis of the other models suggests that the transcription patterns in BM3 are very similar to BM6 and tend to cluster together. The MacTNFa-CR model (**Figure 6B**) predicted potency by classifying donors CT1, CT2, and BM7 as low, medium, and high potency, respectively. The highly potent donors, BM4 and BM5, cluster with BM7. Additionally, the donors BM3 and BM6 correctly cluster around CT2 in the center of the plot. CT3 was incorrectly predicted to have low potency in this model (clustered with CT1), suggesting that the similarity in the transcriptomes of CT3 and CT1 is outweighing the influence of the genes correlated with the potency. CC1 of the CD4Proliferation-OOT model (**Figure 6C**) designates an axis separating BM2, CT2, and BM5 as low, medium, and high potency. In particular, this model contrasts the gene expression between BM2 and BM5. While this model was able predict BM4 and BM7 as low and high potency, respectively, the majority of the other donors clustered centrally with CT2. In the CD4Proliferation-CR model (**Figure 6D**), the main separation was identified between BM and CT MSC donors. While this model predominantly clustered all BM donors together, it was the only model that was able to accurately differentiate between the BM3 and BM6 donors. Notably, a wide distribution represents variability in the genes correlated with the designated metric – for example, in MacTNFa-CR the CT2 sample (**Figure 6B**) shows a bimodal distribution whereas for CD4 proliferation (Figure 5D) there is a single large peak for CT2, suggesting a subpopulation within CT2 donor has more influence on the macrophage TNF alpha response.

## DISCUSSION

A recent surge in the numbers of clinical trials on the use of MSCs for the treatment of various diseases and the management of inflammatory conditions has renewed a lot of interests in understanding MSC-based cell therapy products through deep phenotypic and functional characterization. Due to diverse molecular and secretory functions, MSCs can be used to treat a large number of diseases and immune disorders; however, the disease specific mechanisms of action of MSCs are still unknown (1). Ongoing clinical uses of MSCs rely majorly on their hypothesized immunomodulatory functions, which are to yet be verified in appropriate disease-relevant in vivo models. One of the commonly hypothesized mechanisms of MSC mediated immunomodulation is their ability to suppress T cell upon systemic administration and thus *in vitro* T cell proliferation assay is commonly being used as a potency assay to assess functional properties of MSC cell therapy products (11–13). However, a few recent reports suggest that the macrophages/monocytes could play central roles in causing T cell suppression following intravenous administration of MSCs (14, 16). In this study, we used both macrophage and T cell-based *in vitro* functional assays to compare the immunomodulatory potency of BM and CT-MSCs. Moreover, we compared fresh thaw/out-of-thaw (i.e., immediately after thawing a cryopreserved vial) MSCs with culture-rescue MSCs (cultured for 1-2 days after thawing a cryopreserved vials) in these functional assays. Fresh thaw/out-of-thaw MSCs are often used in the clinics to simplify dose delivery to patient and to avoid the logistic challenges that are associated with the use of freshly harvested or culture-rescued MSCs.

In our macrophage activation assay, both OOT and CR conditions for BM and CT-MSCs significantly suppressed various pro-inflammatory cytokines, including TNFα, IL-12 and MIP-1α. These cytokines are known to be elevated as a result of cytokine storm due to the activation of innate immune cells, including macrophages during acute viral and bacterial infection (e.g., SARS-COV-2 infection (21, 22), sepsis (23, 24)), and BM and CT-MSCs (both OOT and CR products) may mitigate the hyperinflammation by inhibiting the cytokine response of macrophages (25, 26). In T cell proliferation assay, OOT MSCs showed very low or no suppression as compared to CR MSCs for both BM and CT MSCs. Similar differences in T cell proliferation suppression were previously observed between fresh and cryopreserved (out-of-thaw) MSCs (11). T cell proliferation assays can be done using various methods, such as CFSE dilution (13, 27), Ki67 staining (11) and thymidine uptake (28). While all these T cell proliferation assays can quantitatively measure T cell proliferation percentage, the absolute percentage of T cell proliferation (normalized to corresponding activated T cell control) for test groups may differ across these assays. We used CFSE labeled, anti-CD3/CD28 magnetic bead activated PBMCs and co-cultured with MSCs (without irradiation) for 3 days before measuring CSFE dilution in the proliferating CD4 and CD8 T cells. In our flow cytometry-based CSFE-labeled T cell proliferation assay, CR MSCs (for both BM and CT-MSCs) consistently showed suppression of both CD4 and CD8 T cell proliferation in the range of 50-90% with some variabilities across donors. For some BM and CT-MSC donors, OOT MSCs showed up to 30% T cell proliferation suppression.

Additionally, we measured cytokine and chemokine levels in the culture supernatant and found that OOT and CR conditions (from BM and CT-MSCs) showed decreased levels of multiple cytokines and chemokines as compared to the activated PBMC groups with some degree of variability between BM and CT-MSCs and their OOT and CR conditions. Interestingly, some of the cytokines that are known to be secreted by the activated T cells, such as IFN-gamma, IL-17A, IL-2R, TNF-α, IL-5 were inhibited by either OOT or CR conditions for BM or CT MSCs. IFN-gamma cytokine mirrored the T cell proliferation results where CR conditions were more suppressive than OOT conditions across BM and CT MSCs. However, other activated T cell related cytokines, such as IL17A, IL2-R, TNFα, IL5 did not follow the same trend as IFN-gamma. In fact, both OOT and CR conditions from BM and CT-MSCs inhibited at least one of these cytokines. Together, these results suggest differential effect of OOT and CR conditions on activated T cell proliferation and their cytokine secretion.

Next, we performed scRNAseq analysis of out-of-thaw MSCs (as cell therapy product) to predict their immunomodulatory potency (in *in vitro* macrophage activation and T cell proliferation assays) from their transcriptomic attributes. We recently reported the detailed method of scRNAseq analysis and gene expression profiles of out-of-thaw BM and CT-MSCs(17). Furthermore, we showed tissue-source and cell cycle-dependent differential gene expression between BM and CT MSCs and the heterogeneity within each BM or CT MSC donor. In this study, we included a few extra BM-MSC samples expanded either with regular or xeno-free media and used CT-MSC samples from the same donor as before but with one passage later. We did not include a few previous reported BM and CT-MSC donors as we did not have any potency data on those samples. However as before, we observed heterogeneity and tissue source and cell cycle-dependent differential gene expression profile in this cohort of samples. Additionally, we observed culture media-type dependent differential gene expressions for BM-MSCs.

One of our main goals in this study was to predict their immunomodulatory potency of MSCs from their transcriptome profile. For this, we developed two types of predictive models (SR and CCA) using top 1000 differentially expressed genes between OOT BM and CT-MSCs as “inputs” and macrophage TNF alpha (MacTNFa) or CD4 T cell proliferation (CD4Proliferation) response for OOT or CR conditions as separate “output”. SR modeling was used to identify transcriptomic features that predict the potency whereas CCA modeling was used to predict donors with high or low potency.

In the SR models, when we included the tissue source as a potential variable, it came out as the top variable feature correlating with the MacTNFa and CD4Proliferation. In addition, multiple transcriptomic features were also found to be top driving variables for the MacTNFa and CD4Proliferation responses, indicating that these features could potentially be used as quality attributes. However, it is important to note that suitable validation studies using new BM and CT-MSCs samples are needed before these features can be used as critical quality attributes (CQAs). Notably, we did not find any common transcriptomic features that correlate with both MacTNFa and CD4Proliferation responses, which could be due to the differences in the immunomodulatory effects of MSCs in these two potency assays. One of the limitations of our current SR models are the total number of MSC samples (n=10) and number of donors (n=8). The number of donors imposed limits on allowable complexity to avoid overfitting and precluded the inclusion of donors as an input variable.

For CCA, four models were trained using separate output metrics to define MSC potency. Overall, the models trained on MacTNFa data had better predictive capacity than the models trained on CD4 proliferation data. In both MacTNFa models, the predicted order of donor potency followed the recorded *in vitro* potency data, with the exception of a single donor. The CD4 proliferation models were heavily influenced by transcription differences associated with MSC source. However, while the CD4 proliferation models tended to be less capable of predicting the overall order of potency, they were able to identify potency differences in the different BM donors that the MacTNFa models could not. For example, BM2 and BM5 are completely contrasted in Figure 5c and this difference is also apparent in the TNF-α data but is not represented by the MacTNFa models. This highlights the variability in the estimated potency as a function of the test being used to assess potency, as well as the overall similarity in gene expression between donors. There are multiple similarities in the predicted potency of the donors across the four models, despite being trained on different subsets of data and using different potency metrics (i.e., BM3 is always at least 80% overlapped with BM7). One main limitation of CCA as a predictive model is that each component is a linear combination of every input feature without reduction of the feature set. As such, high expression of genes that have minimal correlation with potency can create significant noise.

In summary, we showed here that the OOT and CR conditions perform differently in macrophage activation and T cell proliferation assays. However, BM and CT-MSCs have comparable suppressive effect in these functional assays. Out-of-thaw BM and CT-MSCs differ in their transcriptome profile at single-cell level with some degree of heterogeneity within each donor. Using SR modelling, we have identified multiple transcriptomic features of out-of-thaw MSCs that correlate with their macrophage TNF-α and CD4 T cell proliferation responses of OOT and CR conditions. Also, we have developed CCA models that can predict MSC donors with high or low potency for OOT and CR macrophage TNF alpha and CD4Proliferation responses from their OOT transcriptomic attributes.

## ACKNOWLEDGEMENTS

We are thankful to Wesley Grove for his help in the cell culture and setting up the T cell assay. This work was supported by The Billie and Bernie Marcus Foundation through their generous gift to the Marcus Center for Therapeutic Cell Characterization and Manufacturing (MC3M), the Georgia Research Alliance, and the National Science Foundation Engineering Research Center for Cell Manufacturing Technologies (NSF EEC 1648035).

## AUTHOR CONTRIBUTIONS

PP, CY, KR conceptualized studies; PP, PC, HYS, AJ, LK, WJS, YL performed the experiments, CG developed CCA models; CMT and PC analyzed scRNAseq data; TK and PC developed SR models; PP, PC, HYS, CG, CMT, AJ, LK, YL wrote the manuscript; All authors edited and revised the manuscript; JK, GG, CY, KR provided resources and guidance; CY and KR supervised the studies.

