## Supplemental Information for "Single-Cell Transcriptomic Attributes and Unbiased Computational Modeling for the Prediction of Immunomodulatory Potency of Mesenchymal Stromal Cells"

### <u>SUPPLEMETNAL INFORMATION</u>

**Table S1 List of BM and CT-MSC samples used for omics characterization and potency assays**

| Labels | MSC Donors | Tissue Source | Passage |
|---|---|---|---|
| BM1 | RB139 | BM | P3 |
| BM2 | RB168 (Xeno-Free for RB179) | | |
| BM3 | RB171 (Xeno-Free for RB183) | | |
| BM4 | RB174 | | |
| BM5 | RB177 | | |
| BM6 | RB179 | | |
| BM7 | RB183 | | |
| CT1 | | CT | P3 |
| CT2 | | | |
| CT3 | | | |

**Table S2 BM and CT-MSC donors' information from Single-cell RNA-seq analysis**

| Sample | Number of Cells | Number of Reads | Mean Reads per cell | Median genes per cell | Number of cells post filtering | Sex | Age |
|---|---|---|---|---|---|---|---|
| BM1 (RB139) | 293 | 29,671,530 | 15,469 | 2,218 | 293 | Male | 25 |
| BM2 (RB168) | 359 | 27,494,535 | 37,242 | 4,372 | 333 | Male | 21 |
| BM3 (RB171) | 528 | 33,688,175 | 26,462 | 3,854 | 502 | Female | 26 |
| BM4 (RB174) | 365 | 32,280,924 | 44,038 | 3,763 | 287 | Male | 25 |
| BM5 (RB177) | 394 | 75,876,813 | 101,364 | 4,547 | 355 | Male | 22 |
| BM6 (RB179) | 310 | 29,943,109 | 46,467 | 3,855 | 252 | Male | 21 |
| BM7 (RB183) | 325 | 54,520,104 | 55,728 | 2,861 | 256 | Female | 26 |
| CT1 | 329 | 136,225,518 | 114,420 | 2,733 | 321 | Male | -- |
| CT2 | 141 | 90,952,303 | 91,876 | 1,919 | 141 | Male | -- |
| CT3 | 290 | 81,813,660 | 120,147 | 4,151 | 249 | Male | -- |

**Table S3 List of top variables present in the top performing symbolic regression (SR) models. Yellow highlights indicate if tissue types were allowed as a potential variable. Green highlights indicate the variable was present in >30% of the top performing models**

| ANOVA Trim Results Variables Present in Top Performing Models | | | | | | | | |
|---|---|---|---|---|---|---|---|---|
| **Method** | **OOT** | **OOT** | **CR** | **CR** | **OOT** | **OOT** | **CR** | **CR** |
| **Response** | **TNFa** | **TNFa** | **TNFa** | **TNFa** | **CD4T prol** | **CD4T prol** | **CD4T prol** | **CD4T prol** |
| **Allowed Varible Tissue Type** | **Yes** | **No** | **Yes** | **No** | **Yes** | **No** | **Yes** | **No** |
| **R2** | **91%** | **84%** | **65%** | **63%** | **58%** | **54%** | **48%** | **48%** |
| **Driving Variables** | ***Percent of Variables Present in Top Performming Models*** | | | | | | | |
| **Tissue Type** | 100% | | 100% | | | | 100% | |
| POSTN | 73% | | | | | | | |
| CST3 | 60% | | | | | | | |
| DCN | 50% | | | | | | | |
| TAGLN | 48% | 100% | | | | | | |
| FTL | 42% | | | | | | | |
| FBLN1 | 42% | | | | | | | |
| IFITM2 | 37% | | | | | 86% | 100% | 100% |
| TUBA1B | 32% | 100% | | | | | | |
| ANXA2P3 | | | 95% | 100% | | | 41% | |
| POLCE | | 39% | 90% | 99% | | | | |
| SRP14 | | | 77% | | | | | |
| TIMP1 | | | 67% | 57% | | | | |
| COL1A2 | | | 40% | | | | | |
| IGFBP7 | | 100% | | | | | | |
| COLEC12 | | 100% | | | | | | |
| HTRA1 | | 83% | | | | | | |
| CTHRC1 | | 81% | | 100% | 94% | 96% | | |
| LTBP1 | | 71% | | 69% | | | | |
| SERINC2 | | 60% | | | | | | 100% |
| S100A4 | | 58% | | | | | | |
| LRRC17 | | 37% | | | | | | |
| RPS14P3 | | | | 85% | | | | |
| PXDN | | | | 95% | | | | 98% |
| HLAB | | | | 90% | | | | |
| VEGFA | | | | | 100% | 98% | | |
| HMGA1 | | | | | 98% | 98% | | |
| FDPS | | | | | 79% | 92% | | |
| CYR61 | | | | | 61% | 97% | 44% | |
| DUSP1 | | | | | 59% | 37% | | 94% |
| FLJ44635 | | | | | 43% | 67% | | |
| CTGF | | | | | 31% | | 63% | |
| SNHG5 | | | | | | | 100% | 91% |
| COL12A1 | | | | | | | 37% | |
| EIF3F | | | | | | | 30% | |
| SLC20A1 | | | | | | 52% | | |
| RPS2 | | | | | | 38% | | |
| CLIC1 | | | | | | 35% | | |
| ANPEP | | | | | | | | 99% |
| COLEC12 | | | | | | | | 86% |
| CYR61 | | | | | | | | 64% |
| FBLN5 | | | | | | | | 44% |

#### Table S4 OOT-TNFa Variable Associations Appearing in ≥ 30% of models

##### Two-Way Variable Associations

| oOTTNFA | | | | | | |
|---|---|---|---|---|---|---|
| Rank | Var 1 | Var 2 | Var 1 % | Var 2 % | ⌗ Models | % of Models |
| 1 | tissue•Type | pOSTN | 72.8 | 100.0 | 115 | 72.8 |
| 2 | tissue•Type | cST3 | 59.5 | 100.0 | 94 | 59.5 |
| 3 | tissue•Type | dCN | 50.0 | 100.0 | 79 | 50.0 |
| 4 | cST3 | dCN | 81.9 | 97.5 | 77 | 48.7 |
| 5 | tissue•Type | tAGLN | 48.1 | 100.0 | 76 | 48.1 |
| 6 | tissue•Type | fTL | 41.8 | 100.0 | 66 | 41.8 |
| 7 | tissue•Type | fBLN1 | 41.8 | 100.0 | 66 | 41.8 |
| 8 | cST3 | fTL | 70.2 | 100.0 | 66 | 41.8 |
| 9 | dCN | fTL | 82.3 | 98.5 | 65 | 41.1 |
| 10 | pOSTN | cST3 | 54.8 | 67.0 | 63 | 39.9 |
| 11 | pOSTN | tAGLN | 52.2 | 78.9 | 60 | 38.0 |
| 12 | pOSTN | dCN | 51.3 | 74.7 | 59 | 37.3 |
| 13 | tissue•Type | iFITM2 | 36.7 | 100.0 | 58 | 36.7 |
| 14 | fBLN1 | iFITM2 | 83.3 | 94.8 | 55 | 34.8 |
| 15 | tissue•Type | tUBA1B | 31.6 | 100.0 | 50 | 31.6 |
| 16 | pOSTN | fTL | 42.6 | 74.2 | 49 | 31.0 |
| 17 | pOSTN | fBLN1 | 42.6 | 74.2 | 49 | 31.0 |

##### Three-Way Variable Associations

| oOTTNFA | | | | | | | | |
|---|---|---|---|---|---|---|---|---|
| Rank | Var 1 | Var 2 | Var 3 | Var 1 % | Var 2 % | Var 3 % | ⌗ Models | % of Models |
| 1 | tissue•Type | cST3 | dCN | 48.7 | 81.9 | 97.5 | 77 | 48.7 |
| 2 | tissue•Type | cST3 | fTL | 41.8 | 70.2 | 100.0 | 66 | 41.8 |
| 3 | tissue•Type | dCN | fTL | 41.1 | 82.3 | 98.5 | 65 | 41.1 |
| 4 | cST3 | dCN | fTL | 69.1 | 82.3 | 98.5 | 65 | 41.1 |
| 5 | tissue•Type | pOSTN | cST3 | 39.9 | 54.8 | 67.0 | 63 | 39.9 |
| 6 | tissue•Type | pOSTN | tAGLN | 38.0 | 52.2 | 78.9 | 60 | 38.0 |
| 7 | tissue•Type | pOSTN | dCN | 37.3 | 51.3 | 74.7 | 59 | 37.3 |
| 8 | pOSTN | cST3 | dCN | 49.6 | 60.6 | 72.2 | 57 | 36.1 |
| 9 | tissue•Type | fBLN1 | iFITM2 | 34.8 | 83.3 | 94.8 | 55 | 34.8 |
| 10 | tissue•Type | pOSTN | fTL | 31.0 | 42.6 | 74.2 | 49 | 31.0 |
| 11 | tissue•Type | pOSTN | fBLN1 | 31.0 | 42.6 | 74.2 | 49 | 31.0 |
| 12 | pOSTN | cST3 | fTL | 42.6 | 52.1 | 74.2 | 49 | 31.0 |
| 13 | pOSTN | dCN | fTL | 42.6 | 62.0 | 74.2 | 49 | 31.0 |

##### Four-Way Variable Associations

| oOTTNFA | | | | | | | | | | |
|---|---|---|---|---|---|---|---|---|---|---|
| Rank | Var 1 | Var 2 | Var 3 | Var 4 | Var 1 % | Var 2 % | Var 3 % | Var 4 % | ⌗ Models | % of Models |
| 1 | tissue•Type | cST3 | dCN | fTL | 41.1 | 69.1 | 82.3 | 98.5 | 65 | 41.1 |
| 2 | tissue•Type | pOSTN | cST3 | dCN | 36.1 | 49.6 | 60.6 | 72.2 | 57 | 36.1 |
| 3 | tissue•Type | pOSTN | cST3 | fTL | 31.0 | 42.6 | 52.1 | 74.2 | 49 | 31.0 |
| 4 | tissue•Type | pOSTN | dCN | fTL | 31.0 | 42.6 | 62.0 | 74.2 | 49 | 31.0 |
| 5 | pOSTN | cST3 | dCN | fTL | 42.6 | 52.1 | 62.0 | 74.2 | 49 | 31.0 |

##### Five-Way Variable Association

| oOTTNFA | | | | | | | | | | | | |
|---|---|---|---|---|---|---|---|---|---|---|---|---|
| Rank | Var 1 | Var 2 | Var 3 | Var 4 | Var 5 | Var 1 % | Var 2 % | Var 3 % | Var 4 % | Var 5 % | ⌗ Models | % of Models |
| 1 | tissue•Type | pOSTN | cST3 | dCN | fTL | 31.0 | 42.6 | 52.1 | 62.0 | 74.2 | 49 | 31.0 |

#### Table S5 CR-TNFa Variable Associations Appearing in ≥ 30% of models

##### Two-Way Variable Associations

| cRTNFA | | | | | | |
|---|---|---|---|---|---|---|
| Rank | Var 1 | Var 2 | Var 1 % | Var 2 % | # Models | % of Models |
| 1 | tissue•Type | aNXA2P3 | 95.1 | 100.0 | 234 | 95.1 |
| 2 | tissue•Type | pCOLCE | 90.2 | 100.0 | 222 | 90.2 |
| 3 | aNXA2P3 | pCOLCE | 89.7 | 94.6 | 210 | 85.4 |
| 4 | tissue•Type | sRP14 | 76.8 | 100.0 | 189 | 76.8 |
| 5 | aNXA2P3 | sRP14 | 77.4 | 95.8 | 181 | 73.6 |
| 6 | pCOLCE | sRP14 | 74.8 | 87.8 | 166 | 67.5 |
| 7 | tissue•Type | tIMP1 | 66.7 | 100.0 | 164 | 66.7 |
| 8 | aNXA2P3 | tIMP1 | 67.1 | 95.7 | 157 | 63.8 |
| 9 | sRP14 | tIMP1 | 76.7 | 88.4 | 145 | 58.9 |
| 10 | pCOLCE | tIMP1 | 64.0 | 86.6 | 142 | 57.7 |
| 11 | tissue•Type | cOL1A2 | 40.2 | 100.0 | 99 | 40.2 |
| 12 | aNXA2P3 | cOL1A2 | 41.0 | 97.0 | 96 | 39.0 |
| 13 | pCOLCE | cOL1A2 | 41.0 | 91.9 | 91 | 37.0 |

##### Three-Way Variable Associations

| cRTNFA | | | | | | | | |
|---|---|---|---|---|---|---|---|---|
| Rank | Var 1 | Var 2 | Var 3 | Var 1 % | Var 2 % | Var 3 % | # Models | % of Models |
| 1 | tissue•Type | aNXA2P3 | pCOLCE | 85.4 | 89.7 | 94.6 | 210 | 85.4 |
| 2 | tissue•Type | aNXA2P3 | sRP14 | 73.6 | 77.4 | 95.8 | 181 | 73.6 |
| 3 | tissue•Type | pCOLCE | sRP14 | 67.5 | 74.8 | 87.8 | 166 | 67.5 |
| 4 | aNXA2P3 | pCOLCE | sRP14 | 67.5 | 71.2 | 83.6 | 158 | 64.2 |
| 5 | tissue•Type | aNXA2P3 | tIMP1 | 63.8 | 67.1 | 95.7 | 157 | 63.8 |
| 6 | tissue•Type | sRP14 | tIMP1 | 58.9 | 76.7 | 88.4 | 145 | 58.9 |
| 7 | tissue•Type | pCOLCE | tIMP1 | 57.7 | 64.0 | 86.6 | 142 | 57.7 |
| 8 | aNXA2P3 | sRP14 | tIMP1 | 59.8 | 74.1 | 85.4 | 140 | 56.9 |
| 9 | aNXA2P3 | pCOLCE | tIMP1 | 57.7 | 60.8 | 82.3 | 135 | 54.9 |
| 10 | pCOLCE | sRP14 | tIMP1 | 55.4 | 65.1 | 75.0 | 123 | 50.0 |
| 11 | tissue•Type | aNXA2P3 | cOL1A2 | 39.0 | 41.0 | 97.0 | 96 | 39.0 |
| 12 | tissue•Type | pCOLCE | cOL1A2 | 37.0 | 41.0 | 91.9 | 91 | 37.0 |
| 13 | aNXA2P3 | pCOLCE | cOL1A2 | 37.6 | 39.6 | 88.9 | 88 | 35.8 |

##### Four-Way Variable Associations

| cRTNFA | | | | | | | | | | |
|---|---|---|---|---|---|---|---|---|---|---|
| Rank | Var 1 | Var 2 | Var 3 | Var 4 | Var 1 % | Var 2 % | Var 3 % | Var 4 % | # Models | % of Models |
| 1 | tissue•Type | aNXA2P3 | pCOLCE | sRP14 | 64.2 | 67.5 | 71.2 | 83.6 | 158 | 64.2 |
| 2 | tissue•Type | aNXA2P3 | sRP14 | tIMP1 | 56.9 | 59.8 | 74.1 | 85.4 | 140 | 56.9 |
| 3 | tissue•Type | aNXA2P3 | pCOLCE | tIMP1 | 54.9 | 57.7 | 60.8 | 82.3 | 135 | 54.9 |
| 4 | tissue•Type | pCOLCE | sRP14 | tIMP1 | 50.0 | 55.4 | 65.1 | 75.0 | 123 | 50.0 |
| 5 | aNXA2P3 | pCOLCE | sRP14 | tIMP1 | 50.4 | 53.2 | 62.4 | 72.0 | 118 | 48.0 |
| 6 | tissue•Type | aNXA2P3 | pCOLCE | cOL1A2 | 35.8 | 37.6 | 39.6 | 88.9 | 88 | 35.8 |

##### Five-Way Variable Association

| cRTNFA | | | | | | | | | | | | |
|---|---|---|---|---|---|---|---|---|---|---|---|---|
| Rank | Var 1 | Var 2 | Var 3 | Var 4 | Var 5 | Var 1 % | Var 2 % | Var 3 % | Var 4 % | Var 5 % | # Models | % of Models |
| 1 | tissue•Type | aNXA2P3 | pCOLCE | sRP14 | tIMP1 | 48.0 | 50.4 | 53.2 | 62.4 | 72.0 | 118 | 48.0 |

### Table S6 OOT-CD4T pol Variable Associations Appearing in ≥ 30% of models

#### Two-Way Variable Associations

| oOTCD4T•prol | | | | | | |
|---|---|---|---|---|---|---|
| Rank | Var 1 | Var 2 | Var 1 % | Var 2 % | ⌗ Models | % of Models |
| 1 | vEGFA | hMGA1 | 98.1 | 100.0 | 203 | 98.1 |
| 2 | vEGFA | cTHRC1 | 94.2 | 100.0 | 195 | 94.2 |
| 3 | hMGA1 | cTHRC1 | 94.1 | 97.9 | 191 | 92.3 |
| 4 | vEGFA | fDPS | 79.2 | 100.0 | 164 | 79.2 |
| 5 | hMGA1 | fDPS | 78.8 | 97.6 | 160 | 77.3 |
| 6 | cTHRC1 | fDPS | 80.0 | 95.1 | 156 | 75.4 |
| 7 | vEGFA | cYR61 | 61.4 | 100.0 | 127 | 61.4 |
| 8 | hMGA1 | cYR61 | 60.6 | 96.9 | 123 | 59.4 |
| 9 | vEGFA | dUSP1 | 58.9 | 100.0 | 122 | 58.9 |
| 10 | cTHRC1 | cYR61 | 61.5 | 94.5 | 120 | 58.0 |
| 11 | hMGA1 | dUSP1 | 58.1 | 96.7 | 118 | 57.0 |
| 12 | cTHRC1 | dUSP1 | 59.5 | 95.1 | 116 | 56.0 |
| 13 | fDPS | dUSP1 | 61.0 | 82.0 | 100 | 48.3 |
| 14 | fDPS | cYR61 | 54.9 | 70.9 | 90 | 43.5 |
| 15 | vEGFA | fLJ44635 | 42.5 | 100.0 | 88 | 42.5 |
| 16 | hMGA1 | fLJ44635 | 43.3 | 100.0 | 88 | 42.5 |
| 17 | cTHRC1 | fLJ44635 | 39.5 | 87.5 | 77 | 37.2 |
| 18 | cYR61 | dUSP1 | 54.3 | 56.6 | 69 | 33.3 |
| 19 | vEGFA | cTGF | 31.4 | 100.0 | 65 | 31.4 |
| 20 | hMGA1 | cTGF | 32.0 | 100.0 | 65 | 31.4 |
| 21 | fDPS | cTGF | 39.0 | 98.5 | 64 | 30.9 |
| 22 | cTHRC1 | cTGF | 32.3 | 96.9 | 63 | 30.4 |

#### Three-Way Variable Associations

| oOTCD4T•prol | | | | | | | | |
|---|---|---|---|---|---|---|---|---|
| Rank | Var 1 | Var 2 | Var 3 | Var 1 % | Var 2 % | Var 3 % | ⌗ Models | % of Models |
| 1 | vEGFA | hMGA1 | cTHRC1 | 92.3 | 94.1 | 97.9 | 191 | 92.3 |
| 2 | vEGFA | hMGA1 | fDPS | 77.3 | 78.8 | 97.6 | 160 | 77.3 |
| 3 | vEGFA | cTHRC1 | fDPS | 75.4 | 80.0 | 95.1 | 156 | 75.4 |
| 4 | hMGA1 | cTHRC1 | fDPS | 74.9 | 77.9 | 92.7 | 152 | 73.4 |
| 5 | vEGFA | hMGA1 | cYR61 | 59.4 | 60.6 | 96.9 | 123 | 59.4 |
| 6 | vEGFA | cTHRC1 | cYR61 | 58.0 | 61.5 | 94.5 | 120 | 58.0 |
| 7 | vEGFA | hMGA1 | dUSP1 | 57.0 | 58.1 | 96.7 | 118 | 57.0 |
| 8 | vEGFA | cTHRC1 | dUSP1 | 56.0 | 59.5 | 95.1 | 116 | 56.0 |
| 9 | hMGA1 | cTHRC1 | cYR61 | 57.1 | 59.5 | 91.3 | 116 | 56.0 |
| 10 | hMGA1 | cTHRC1 | dUSP1 | 55.2 | 57.4 | 91.8 | 112 | 54.1 |
| 11 | vEGFA | fDPS | dUSP1 | 48.3 | 61.0 | 82.0 | 100 | 48.3 |
| 12 | cTHRC1 | fDPS | dUSP1 | 50.3 | 59.8 | 80.3 | 98 | 47.3 |
| 13 | hMGA1 | fDPS | dUSP1 | 47.3 | 58.5 | 78.7 | 96 | 46.4 |
| 14 | vEGFA | fDPS | cYR61 | 43.5 | 54.9 | 70.9 | 90 | 43.5 |
| 15 | vEGFA | hMGA1 | fLJ44635 | 42.5 | 43.3 | 100.0 | 88 | 42.5 |
| 16 | hMGA1 | fDPS | cYR61 | 42.4 | 52.4 | 67.7 | 86 | 41.5 |
| 17 | cTHRC1 | fDPS | cYR61 | 43.6 | 51.8 | 66.9 | 85 | 41.1 |
| 18 | vEGFA | cTHRC1 | fLJ44635 | 37.2 | 39.5 | 87.5 | 77 | 37.2 |
| 19 | hMGA1 | cTHRC1 | fLJ44635 | 37.9 | 39.5 | 87.5 | 77 | 37.2 |
| 20 | vEGFA | cYR61 | dUSP1 | 33.3 | 54.3 | 56.6 | 69 | 33.3 |
| 21 | cTHRC1 | cYR61 | dUSP1 | 33.8 | 52.0 | 54.1 | 66 | 31.9 |
| 22 | vEGFA | hMGA1 | cTGF | 31.4 | 32.0 | 100.0 | 65 | 31.4 |
| 23 | hMGA1 | cYR61 | dUSP1 | 32.0 | 51.2 | 53.3 | 65 | 31.4 |
| 24 | vEGFA | fDPS | cTGF | 30.9 | 39.0 | 98.5 | 64 | 30.9 |
| 25 | hMGA1 | fDPS | cTGF | 31.5 | 39.0 | 98.5 | 64 | 30.9 |
| 26 | vEGFA | cTHRC1 | cTGF | 30.4 | 32.3 | 96.9 | 63 | 30.4 |
| 27 | hMGA1 | cTHRC1 | cTGF | 31.0 | 32.3 | 96.9 | 63 | 30.4 |

#### Four-Way Variable Associations

| oOTCD4T•prol | | | | | | | | | | |
|---|---|---|---|---|---|---|---|---|---|---|
| Rank | Var 1 | Var 2 | Var 3 | Var 4 | Var 1 % | Var 2 % | Var 3 % | Var 4 % | ⌗ Models | % of Models |
| 1 | vEGFA | hMGA1 | cTHRC1 | fDPS | 73.4 | 74.9 | 77.9 | 92.7 | 152 | 73.4 |
| 2 | vEGFA | hMGA1 | cTHRC1 | cYR61 | 56.0 | 57.1 | 59.5 | 91.3 | 116 | 56.0 |
| 3 | vEGFA | hMGA1 | cTHRC1 | dUSP1 | 54.1 | 55.2 | 57.4 | 91.8 | 112 | 54.1 |
| 4 | vEGFA | cTHRC1 | fDPS | dUSP1 | 47.3 | 50.3 | 59.8 | 80.3 | 98 | 47.3 |
| 5 | vEGFA | hMGA1 | fDPS | dUSP1 | 46.4 | 47.3 | 58.5 | 78.7 | 96 | 46.4 |
| 6 | hMGA1 | cTHRC1 | fDPS | dUSP1 | 46.3 | 48.2 | 57.3 | 77.0 | 94 | 45.4 |
| 7 | vEGFA | hMGA1 | fDPS | cYR61 | 41.5 | 42.4 | 52.4 | 67.7 | 86 | 41.5 |
| 8 | vEGFA | cTHRC1 | fDPS | cYR61 | 41.1 | 43.6 | 51.8 | 66.9 | 85 | 41.1 |
| 9 | hMGA1 | cTHRC1 | fDPS | cYR61 | 39.9 | 41.5 | 49.4 | 63.8 | 81 | 39.1 |
| 10 | vEGFA | hMGA1 | cTHRC1 | fLJ44635 | 37.2 | 37.9 | 39.5 | 87.5 | 77 | 37.2 |
| 11 | vEGFA | cTHRC1 | cYR61 | dUSP1 | 31.9 | 33.8 | 52.0 | 54.1 | 66 | 31.9 |
| 12 | vEGFA | hMGA1 | cYR61 | dUSP1 | 31.4 | 32.0 | 51.2 | 53.3 | 65 | 31.4 |
| 13 | vEGFA | hMGA1 | fDPS | cTGF | 30.9 | 31.5 | 39.0 | 98.5 | 64 | 30.9 |
| 14 | vEGFA | hMGA1 | cTHRC1 | cTGF | 30.4 | 31.0 | 32.3 | 96.9 | 63 | 30.4 |

#### Five-Way Variable Associations

| oOTCD4T•prol | | | | | | | | | | | | |
|---|---|---|---|---|---|---|---|---|---|---|---|---|
| Rank | Var 1 | Var 2 | Var 3 | Var 4 | Var 5 | Var 1 % | Var 2 % | Var 3 % | Var 4 % | Var 5 % | ⌗ Models | % of Models |
| 1 | vEGFA | hMGA1 | cTHRC1 | fDPS | dUSP1 | 45.4 | 46.3 | 48.2 | 57.3 | 77.0 | 94 | 45.4 |
| 2 | vEGFA | hMGA1 | cTHRC1 | fDPS | cYR61 | 39.1 | 39.9 | 41.5 | 49.4 | 63.8 | 81 | 39.1 |

#### Table S7 CR-CD4T pol Variable Associations Appearing in ≥ 30% of models

##### Two-Way Variable Associations

| cRCD4T•prol | | | | | | |
|---|---|---|---|---|---|---|
| Rank | Var 1 | Var 2 | Var 1 % | Var 2 % | ⌗ Models | % of Models |
| 1 | sNHG5 | tissue•Type | 100.0 | 100.0 | 248 | 100.0 |
| 2 | iFITM2 | tissue•Type | 100.0 | 100.0 | 248 | 100.0 |
| 3 | iFITM2 | sNHG5 | 100.0 | 100.0 | 248 | 100.0 |
| 4 | tissue•Type | cTGF | 63.3 | 100.0 | 157 | 63.3 |
| 5 | sNHG5 | cTGF | 63.3 | 100.0 | 157 | 63.3 |
| 6 | iFITM2 | cTGF | 63.3 | 100.0 | 157 | 63.3 |
| 7 | tissue•Type | cYR61 | 43.5 | 100.0 | 108 | 43.5 |
| 8 | sNHG5 | cYR61 | 43.5 | 100.0 | 108 | 43.5 |
| 9 | iFITM2 | cYR61 | 43.5 | 100.0 | 108 | 43.5 |
| 10 | tissue•Type | aNXA2P3 | 41.1 | 100.0 | 102 | 41.1 |
| 11 | sNHG5 | aNXA2P3 | 41.1 | 100.0 | 102 | 41.1 |
| 12 | iFITM2 | aNXA2P3 | 41.1 | 100.0 | 102 | 41.1 |
| 13 | tissue•Type | cOL12A1 | 36.7 | 100.0 | 91 | 36.7 |
| 14 | iFITM2 | cOL12A1 | 36.7 | 100.0 | 91 | 36.7 |
| 15 | sNHG5 | cOL12A1 | 36.7 | 100.0 | 91 | 36.7 |
| 16 | tissue•Type | eIF3F | 30.2 | 100.0 | 75 | 30.2 |
| 17 | iFITM2 | eIF3F | 30.2 | 100.0 | 75 | 30.2 |
| 18 | sNHG5 | eIF3F | 30.2 | 100.0 | 75 | 30.2 |

##### Three-Way Variable Associations

| cRCD4T•prol | | | | | | | | |
|---|---|---|---|---|---|---|---|---|
| Rank | Var 1 | Var 2 | Var 3 | Var 1 % | Var 2 % | Var 3 % | ⌗ Models | % of Models |
| 1 | iFITM2 | sNHG5 | tissue•Type | 100.0 | 100.0 | 100.0 | 248 | 100.0 |
| 2 | sNHG5 | tissue•Type | cTGF | 63.3 | 63.3 | 100.0 | 157 | 63.3 |
| 3 | iFITM2 | tissue•Type | cTGF | 63.3 | 63.3 | 100.0 | 157 | 63.3 |
| 4 | iFITM2 | sNHG5 | cTGF | 63.3 | 63.3 | 100.0 | 157 | 63.3 |
| 5 | sNHG5 | tissue•Type | cYR61 | 43.5 | 43.5 | 100.0 | 108 | 43.5 |
| 6 | iFITM2 | tissue•Type | cYR61 | 43.5 | 43.5 | 100.0 | 108 | 43.5 |
| 7 | iFITM2 | sNHG5 | cYR61 | 43.5 | 43.5 | 100.0 | 108 | 43.5 |
| 8 | sNHG5 | tissue•Type | aNXA2P3 | 41.1 | 41.1 | 100.0 | 102 | 41.1 |
| 9 | iFITM2 | tissue•Type | aNXA2P3 | 41.1 | 41.1 | 100.0 | 102 | 41.1 |
| 10 | iFITM2 | sNHG5 | aNXA2P3 | 41.1 | 41.1 | 100.0 | 102 | 41.1 |
| 11 | iFITM2 | tissue•Type | cOL12A1 | 36.7 | 36.7 | 100.0 | 91 | 36.7 |
| 12 | sNHG5 | tissue•Type | cOL12A1 | 36.7 | 36.7 | 100.0 | 91 | 36.7 |
| 13 | iFITM2 | sNHG5 | cOL12A1 | 36.7 | 36.7 | 100.0 | 91 | 36.7 |
| 14 | iFITM2 | tissue•Type | eIF3F | 30.2 | 30.2 | 100.0 | 75 | 30.2 |
| 15 | sNHG5 | tissue•Type | eIF3F | 30.2 | 30.2 | 100.0 | 75 | 30.2 |
| 16 | iFITM2 | sNHG5 | eIF3F | 30.2 | 30.2 | 100.0 | 75 | 30.2 |

##### Four-Way Variable Associations

| cRCD4T•prol | | | | | | | | | | |
|---|---|---|---|---|---|---|---|---|---|---|
| Rank | Var 1 | Var 2 | Var 3 | Var 4 | Var 1 % | Var 2 % | Var 3 % | Var 4 % | ⌗ Models | % of Models |
| 1 | iFITM2 | sNHG5 | tissue•Type | cTGF | 63.3 | 63.3 | 63.3 | 100.0 | 157 | 63.3 |
| 2 | iFITM2 | sNHG5 | tissue•Type | cYR61 | 43.5 | 43.5 | 43.5 | 100.0 | 108 | 43.5 |
| 3 | iFITM2 | sNHG5 | tissue•Type | aNXA2P3 | 41.1 | 41.1 | 41.1 | 100.0 | 102 | 41.1 |
| 4 | iFITM2 | sNHG5 | tissue•Type | cOL12A1 | 36.7 | 36.7 | 36.7 | 100.0 | 91 | 36.7 |
| 5 | iFITM2 | sNHG5 | tissue•Type | eIF3F | 30.2 | 30.2 | 30.2 | 100.0 | 75 | 30.2 |

A

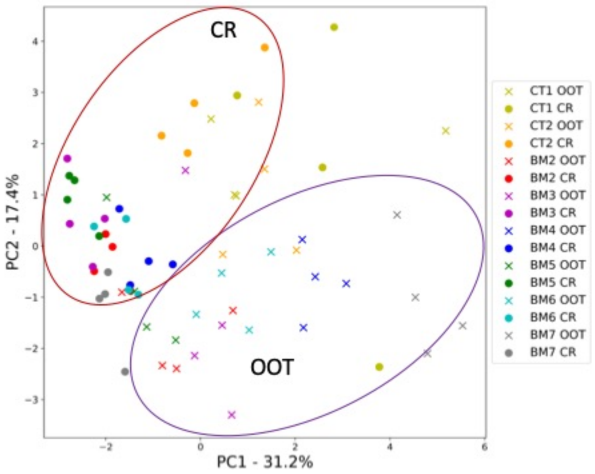


B

**PCA loading scores**

| PC Loadings | With THP-1 M | | Without THP-1 M | |
|---|---|---|---|---|
| | PC1 | PC2 | PC1 | PC2 |
| TNFα | 0.82459 | -0.26281 | 0.0321 | 0.22602 |
| IL12 | 0.86696 | 0.02556 | 0.81577 | -0.16619 |
| MIP1α | 0.65851 | -0.09532 | 0.67081 | 0.52528 |
| MIP1β | 0.6968 | 0.31359 | 0.78435 | 0.20434 |
| MCP-1 | 0.64543 | 0.44288 | 0.80624 | 0.35495 |
| IL1RA | 0.77415 | 0.44425 | 0.89162 | 0.07376 |
| MIG | 0.50987 | 0.71547 | 0.77498 | -0.32128 |
| GCSF | -0.38359 | 0.05354 | -0.10014 | 0.16988 |
| GMCSF | -0.35931 | -0.08099 | -0.07977 | 0.56415 |
| VEGF | -0.64026 | 0.09999 | -0.33061 | -0.36044 |
| RANTES | 0.04409 | 0.11253 | 0.18734 | 0.34484 |
| HGF | -0.10528 | 0.79214 | 0.68252 | -0.1801 |
| IL8 | -0.60665 | 0.44978 | 0.05585 | -0.6816 |
| IL17A | -0.49849 | 0.5433 | 0.16566 | -0.78309 |
| IL13 | -0.36396 | 0.76426 | 0.56509 | -0.56327 |

C

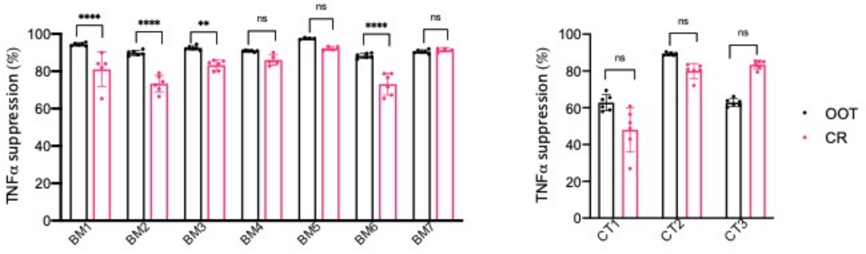


D

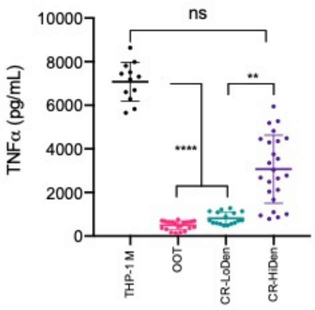


Figure S1. A) Principal component analysis of cytokines, chemokines, and growth factors for OOT and CR conditions (without THP1 macrophage group) for BM and CT-MSCs from macrophage activation assay; B) PCA loading scores of PC1 and PC2 for Figure 2C and Figure S1A; C) Comparative TNF-alpha suppression levels for BM and CT-MSCs from THP-1 macrophage assay; For BM groups, one-way ANOVA with Tukey's post-hoc test was performed. For CT groups, Kruskal-Wallis test with Dunn's post-hoc test was performed. D) Effect of seeding density on TNF alpha response for culture rescue conditions in THP-1 macrophage assay; Kruskal-Wallis test with Dunn's post-hoc test was performed. Data represent mean ±SD. **P<0.01. ***P < 0.001, ****P<0.0001, ns- not significant.

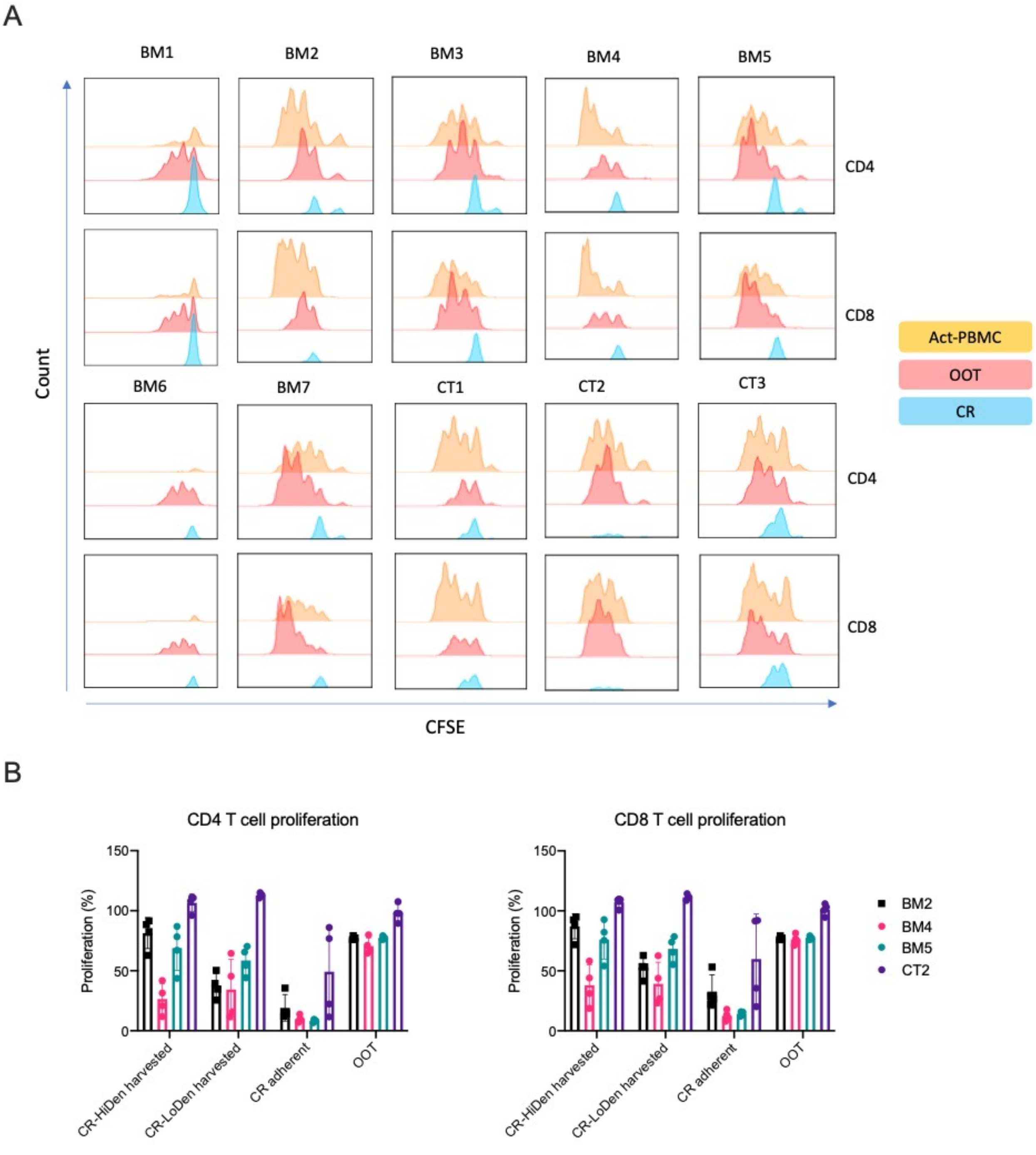


Figure S2. A) Flow histograms showing CFSE dilution due to T cell proliferation for OOT and CR conditions for BM and CT-MSC samples; B) Effect of seeding density of BM and CT-MSCs on T cell proliferation for harvested culture-rescue conditions. Percentage of CD4 and C8 T cell proliferation was normalized to activated PBMC group with 100% proliferation. Data represent mean ±SD.

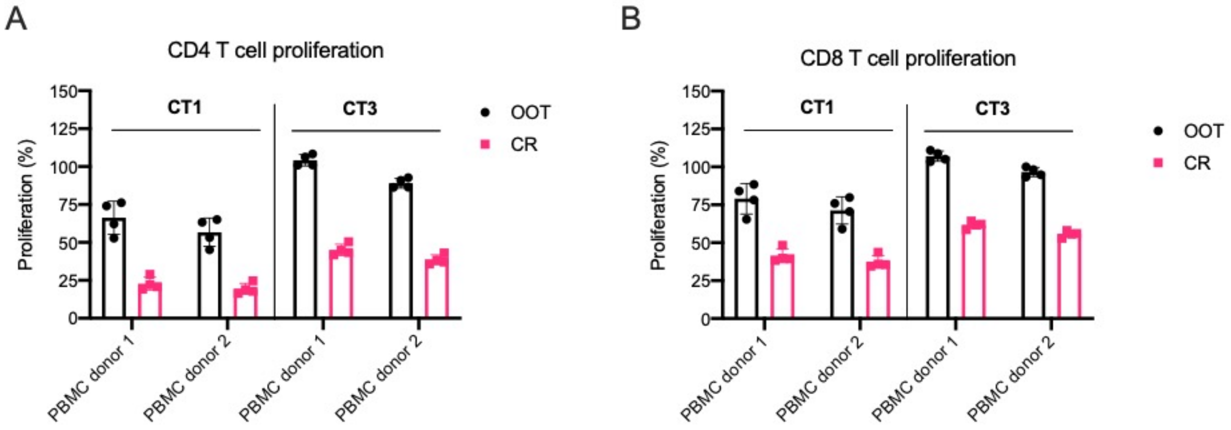


Figure S3. CT1 and CT3 MSCs show similar levels of A) CD4 and B) CD8 T cell proliferation for OOT and CR conditions across two PBMC donors. Percentage of CD4 and C8 T cell proliferation was normalized to activated PBMC group with 100% proliferation. Data represent mean ±SD.

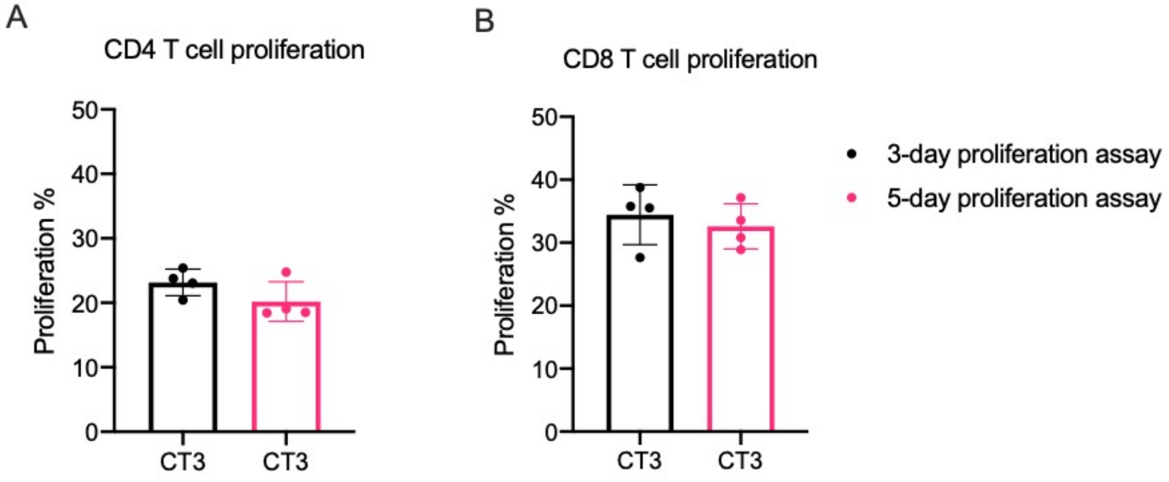


Figure S4. CT3 MSCs show similar levels of A) CD4 and B) CD8 T cell proliferation for CR conditions between 3-day and 5-day proliferation assays. Percentage of CD4 and C8 T cell proliferation was normalized to activated PBMC group with 100% proliferation. Data represent mean ±SD.
